# A distributed lattice of aligned atoms exists in a protein structure: A hierarchical clustering study of displacement parameters in bovine trypsin

**DOI:** 10.1101/475889

**Authors:** Viktor Ahlberg Gagnér, Ida Lundholm, Maria-Jose Garcia-Bonete, Helena Rodilla, Ran Friedman, Vitali Zhaunerchyk, Gleb Bourenkov, Thomas Schneider, Jan Stake, Gergely Katona

## Abstract

Low-frequency vibrations are crucial for protein structure and function, but only a few experimental techniques can shine light on them. The main challenge when addressing protein dynamics in the terahertz domain is the ubiquitous water that exhibit strong absorption. In this paper, we observe the protein atoms directly using X-ray crystallography in bovine trypsin at 100 K while irradiating the crystals with 0.5 THz radiation alternating on and off states. We observed that the anisotropy of atomic displacements increases upon terahertz irradiation. Atomic displacement similarities develop between chemically related atoms and between atoms of the catalytic machinery. This pattern likely arise from delocalized polar vibrational modes rather than delocalized elastic deformations or rigid-body displacements. This method can ultimately reveal how the alignment of chemically related atoms and the underlying polar vibrational dynamics make a protein structure stable.

## Introduction

Enzymes are thermally activated indicating the crucial role of vibrations in their function. Trypsins catalyze the cleavage of peptide bonds in the gut and their catalytic activity also increases with temperature (up to an optimum temperature).(*1*) Trypsin is a globular protein and its folded structure is maintained in a wide range of aqueous solutions. This structure forms spontaneously below a melting temperature, but the conversion from the inactive trypsinogen to the active trypsin form requires the cleavage of the Lys-15/Ile-16 peptide bond by enteropeptidase or an active trypsin molecule. Premature activation of trypsinogen in the acinar cells or the pancreatic duct leads to tissue damage, which contributes to the development of pancreatitis. Through thermal excitation, low-frequency vibrations contribute to folding, conversion between the inactive and active enzyme form (*2*) and the catalytic activity of trypsin. The low-frequency vibrational modes can follow the canonical energy distribution, and such an energy distribution is necessary for most models of enzyme reaction kinetics: the Eyring equation and transition-state theory.(*3*) The ideal behavior is usually only observed in a narrow temperature range in enzyme catalyzed reactions.(*4*) Fröhlich (*5*) predicted an alternative to the canonical energy distribution, which may be more appropriate for open biological dissipative systems (commonly referred to as Fröhlich condensation). In this out-of-equilibrium model, the energy distribution is skewed towards the lowest frequency mode in a coupled oscillator system. Different degrees of condensation have been investigated using simulations and the energy distribution was show to depend on the rates of excitation, the energy exchange between the vibration modes, thermalization and the temperature of the environment. (*6-9*) Collective modes in biological systems are postulated to play a role in long-range protein-protein interactions, the adjustment of biochemical reaction rates and information processing in semicrystalline, supramolecular systems. (*6, 10*) Recently, structural changes were observed in lysozyme crystals when pumped with 0.4 THz radiation at room temperature, which suggests an experimental means to study such phenomena. (*11*) Continuous excitation of fluorophore-labelled bovine serum albumin with visible light has resulted in a terahertz spectral signal, which evolved over minutes time scales. (*12*)

Dynamics also plays a role in long-range allosteric effects as shown by NMR experiments. (*13-15*) It is frequently observed that a mutation or binding of a small molecule at one site can affect the dynamics at remote sites. Computational studies capture some aspects of correlated motions in proteins, (*16, 17*) but experimental benchmarks are often not strict enough for facilitating further improvements. As a general trend, long-range allostery is modeled as a network of interactions from one point to another where the diffusive motions stepwise transmitted through a chain of local interactions between the distant sites. (*18-20*) Local interactions still dominate in these network models and hence the influence they exert is expected to decay with increasing distance as it propagates in space. Reactive interactions of atoms with the alternating electric field are not typically assumed necessary for explaining allosteric changes. Nevertheless, electric field pulses can cause reversible, concerted conformational changes and symmetry breaking in protein crystals.(*21*) Slow, high-field electric pulses do happen infrequently in biological systems near membranes, but terahertz fluctuation are more common in aqueous solutions in the form of thermal vibrations.

The main methods to study low-frequency vibrations in proteins is terahertz spectroscopy, optical Kerr spectroscopy and inelastic/quasielastic neutron scattering. Terahertz spectroscopy (*22*) was particularly successful in separating protein vibrations from the bulk solvent contribution and detected long-range vibrations in the protein scaffold. (*23, 24*) Terahertz spectroscopic measurements also revealed long-range dynamics linked to the functional state of proteins. (*25-27*) The vibrational spectra lack resonant bands and a few polynomials are sufficient to describe the entire terahertz region. Polarization varying anisotropic terahertz microscopy breaks this deadlock by recording two dimensional spectra from oriented proteins in crystals yielding dozens of absorption peaks.(*28*) However, typical proteins contain thousands of atoms and many degrees of freedom. The interpretation of the spectra critically depends on the assumed model of protein and solvent dynamics. High-resolution X-ray crystallography recovers the three dimensional distribution of atomic displacements described with amplitudes and directions.(*29*) If the diffraction experiment is not time-resolved, the displacements are time averaged. They are indistinguishable from a static distribution, but they have a dynamic component. This is a small increase in ambiguity in interpretation, which is compensated by the spatial distribution of every atom in the protein resulting in several thousand parameters. In this study, we demonstrate the use of high-resolution X-ray crystallography to study the terahertz dynamics of proteins. The dynamic component of the distribution was directly investigated by irradiating the crystals with terahertz radiation and compare spatial distributions to the dark state.

The amplitude and direction of atomic motions are valuable constraints for dynamic models. However, the complexity of protein crystal structures makes it difficult to comprehend the patterns that may correspond to the different dynamical models. We solve this problem by applying an unsupervised machine learning technique: hierarchical clustering. The basis of our analysis is the six atomic displacement parameters (ADPs), which can be illustrated with thermal ellipsoids after diagonalization (Figure 1). The axis orientation of the ellipsoids represent the eigenvectors of the ADPs and the amplitudes of the axes correspond to the eigenvalues. For a typical protein atom, the amplitudes are not equal (the atom is anisotropic). To simplify the comparison of the ADPs, we adapted the B_eq_ (equivalent to an isotropic B-factors) (*30*) and anisotropy metrics (*29*) (henceforth ANISO). ANISO varies between zero and one, where one indicates perfect isotropy.

**Figure 1.**
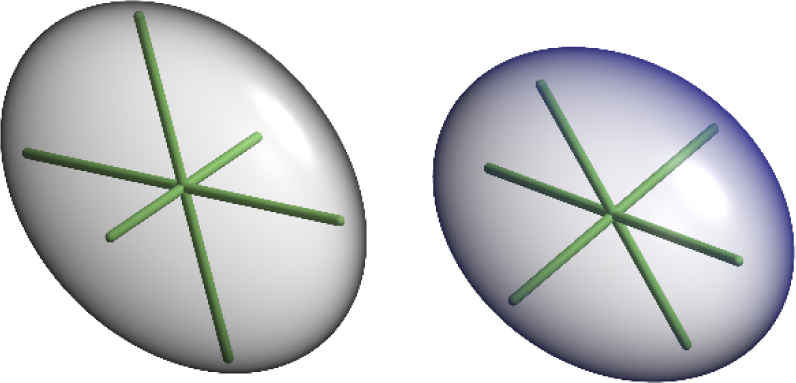
Thermal ellipsoids of the carbonyl carbon (left) and amide nitrogen (right) atoms of Thr-125B located close to each other. The carbonyl carbon and amide nitrogen atoms display slightly oblate (pancake shape) and prolate (cigar shape) ellipsoids, respectively. The direction and length of the axes in the ellipsoids are uniquely defined by eigenvectors and the corresponding eigenvalues of the ADP tensors. These two atoms are located in close vicinity to one other, but they display the highest degree of similarity with distant atoms instead. The figure was obtained using RASTEP (*29*), numbering of amino acid residues follows the chymotrypsin convention. (*31*)

In addition to the traditional directionless metrics (B_eq_ and ANISO), we applied a metric-free approach to include eigenvector directionality in the analysis. The inclusion of all the information from the ADPs dramatically improved the specificity of the clustering. Groups of atoms that share similar dynamics could be inferred from the similarity of their positional probability distributions. The categorization of ADP changes was not only difficult because of the large number of atoms in a protein, but we also had to overcome conceptual challenges. Firstly, going from an U_ij_ matrix to a metric (such as B_eq_ and ANISO) inevitably involves a loss of information (converting six parameters to one). Secondly, it is unlikely that the changes affect only one metric (B_eq_ or ANISO). It is difficult to develop metrics that complement each other well. B_eq_ and ANISO form a reasonable combination, but they do not reveal anything about the directionality of thermal ellipsoids. Thirdly, even the initial diagonalization is problematic. For example, in (partially) isotropic atoms, the eigenvectors become arbitrary and their changes become meaningless and in practice are only influenced by errors. Fourthly, categorizing atoms based on which amino acid type they belong to can be considered artificial as well. Other distinctions may make more sense such as separating main chain from side chain atoms or grouping based on secondary structures, functional groups (carboxyl, hydroxyl, aromatic), atom types etc.

In this study, the non-thermal effect of 0.5 THz radiation was investigated in bovine trypsin crystals cryo-cooled to 100 K. Optimization of the experimental geometry allowed us to collect wider diffraction angles. We analyze the anisotropic displacements in the protein structure at different length scales using different computational methods. We move stepwise from a global overview to atomic details. Initially, we discuss the effects of terahertz irradiation on individual amino residues by averaging the atomic data and comparing those results to simulated data. Finally, we elaborate on the effects on individual atoms related to terahertz dynamics using a clustering method.

## Results

We irradiated orthorhombic bovine trypsin crystals alternately with a 0.5 THz radiation, such that the odd frames correspond to “THz on”-states and the even frames correspond to “THz off”-states. As a reference, we compared them with X-ray diffraction data collected from crystals that were not exposed to terahertz radiation. In contrast to an earlier experiment with hen egg white lysozyme,(*11*) bovine trypsin crystals were cryo-cooled to 100 K during the entire period of data collection with a dry nitrogen gas stream and suspended in a mylar loop without a surrounding plastic capillary. In total, five terahertz irradiated and four reference crystals were retained for analysis. Data processing statistics are summarized in Table S1.

### Comparison of the irradiated and non-irradiated state of the crystals reveal differences between individual residues

An initial comparison of the ADPs was based on the difference in B_eq_-factor between equivalent atoms determined from the odd and even frames (B_eq,odd_-B_eq,even_). The mean and confidence interval (95%) of the difference in B_eq_-factor for amino acid atoms were determined. Figure 2A shows this type of comparison for the twenty amino acids in the reference (*blue*) and terahertz irradiated crystals (*green*). B_eq_ appears to be either unchanged or similar to the references in the terahertz experiments except for the amino acid residue glutamine. The B_eq_ of glycine, glutamine, serine, leucine and methionine atoms appear to decrease in odd data sets in the reference experiments.

**Figure 2.**
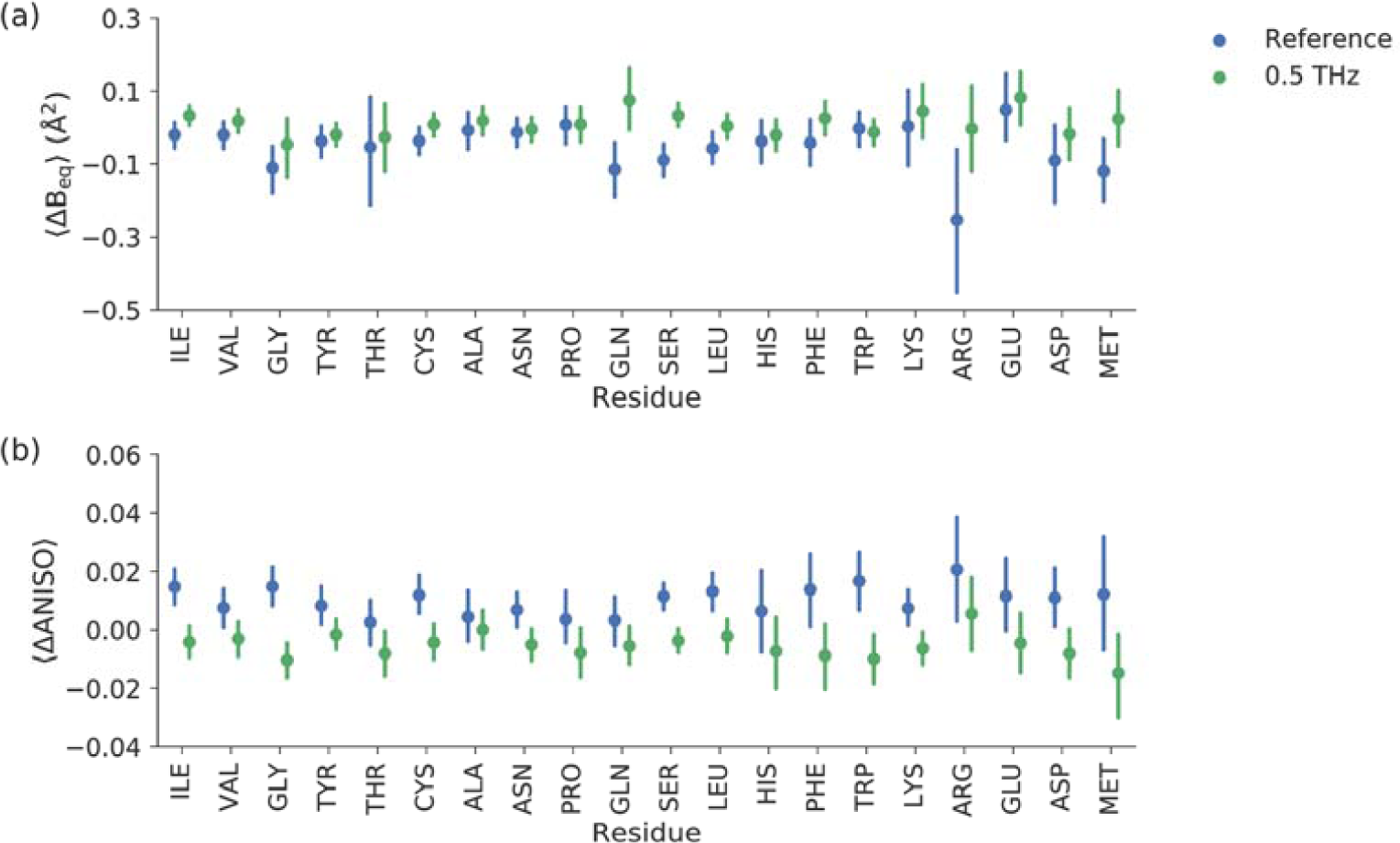
B_eq_-factor and ANISO comparison between odd and even frames of the reference (*blue*) and terahertz irradiated (*green*) crystals in different types of amino acid residues. Each average is calculated from the difference in B_eq_ and ANISO for each individual atom in the reference and the terahertz irradiated crystals, respectively. The error bars represent 95 % CI of the mean.

The comparison of difference ANISO values (Figure 2B) shows a limited overlap of confidence intervals (CI) between the reference and the terahertz categories. The greatest contrast between the reference and terahertz categories are observed in atoms from isoleucine, glycine, cysteine, serine, leucine, tryptophane and lysine residues. In all these cases, the average ANISO of atoms decreases upon terahertz irradiation. The position of the amino acid residue in the structure is important, therefore the ANISO value distribution of individual glycines, tryptophans and histidines are shown in Figure 3. From this comparison, it is obvious that most of the glycine residues behaved similarly in the terahertz radiation experiments compared to the references (Gly-43 is a clear exception), although in general, the atoms of the glycines in the N-terminal domain had a tendency to become more anisotropic when the crystals were subjected to terahertz radiation. Tryptophan residues became more anisotropic when all atoms were pooled together and Trp-51, Trp-215 and Trp-237 had different average anisotropies in relation to the reference, whereas Trp-141 stayed similar. Histidine residues, except catalytic His-57, appeared to be completely unaffected by terahertz radiation in the pooled comparison. As a general tendency, when there was a reversible change in anisotropy upon terahertz irradiation, it predominantly resulted in more anisotropic behavior in the residue atoms.

**Figure 3.**
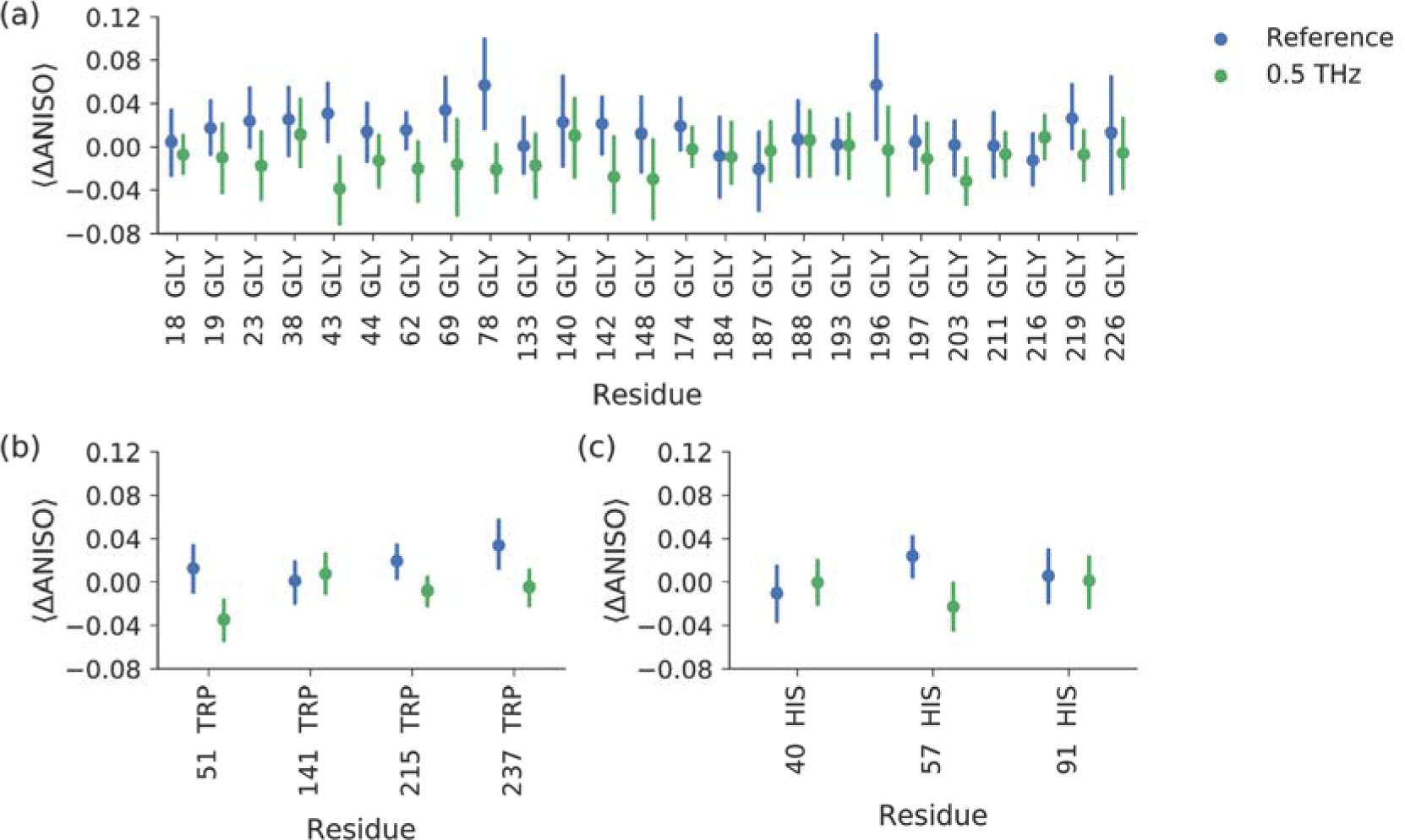
Comparison of ANISO change in selected amino acids (glycines, histidines and tryptophans) between odd and even frames of the reference (*blue*) and terahertz irradiated (*green*) crystals.

### Comparison to NMA and MD simulations

In order to model ADP tensor components (U_ij_) we performed NMA on an energy-minimized trypsin structure and generated random samples of the structure using all normal modes. The calculation assumed a temperature of 100 K. The B_eq_s of predicted atomic U_ij_s were approximately two orders of magnitude smaller than the experimentally observed values. Since NMA only explores the potential surface around a single minimum, it has a limited ability to follow protein dynamics, which consist of multiple conformations that exchange at slower than picoseconds timescales. Moreover, NMA is performed in a vacuum, which is not the case with solvated forms of proteins. Nevertheless, NMA is useful for inferring protein motions and hence we examined the correlation of NMA predicted U_ij_ metrics with Molecular Dynamics (MD) simulations and also with the experimental data.

An alternative to NMA is MD simulations of solvated protein molecules. In order to facilitate conformational changes, we performed the simulations at 300 K. When flash-cooling proteins, conformational variation is locked into a static distribution, but still affects the displacement parameters. Since conformational sampling is more delicate than sampling from an NMA ensemble, we used four MD simulations to predict the B_eq_ and anisotropy of the protein atoms and compared the values obtained to the corresponding parameters of the refined crystal structure (Figure 4). The predicted B_eq_ values were approximately on the same scale as the experimental values. They appeared to be underestimated for the well-ordered parts of the structure, but overestimated for the flexible regions. In particular, the B_eq_ of Tyr, Asn, Gln, Phe and Lys residues were greatly overestimated compared to the experimental data. There was a 0.53 Pearson correlation between the calculated (from segments of four times 10 ns simulations) and the experimental means of B_eq_ per residue (calculated for all equivalent atoms from the reference crystals). The mean ANISO values in amino acid residues showed a smaller correlation (0.39), and MD simulations generally overestimated the anisotropy. By contrast, ANISO values derived from NMA simulations were more isotropic and agreed better with the experimental data. MD simulations predicted Asn residues to be much more anisotropic relative to other amino acid types as well (Figure 4D).

**Figure 4.**
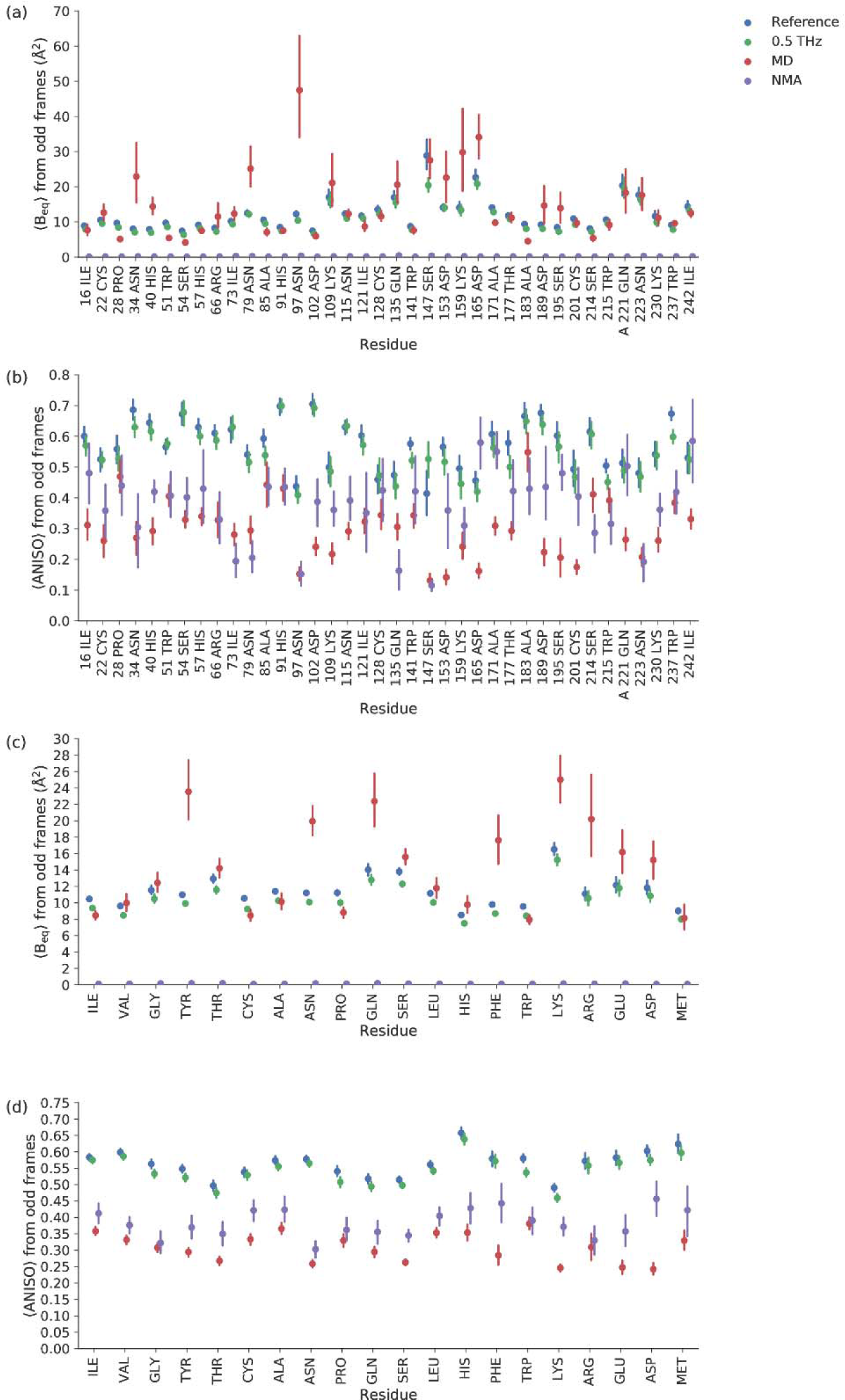
B_eq_-factor and ANISO comparison based on the odd diffraction frames of reference crystals (*blue*) and odd diffraction frames of terahertz irradiated (*green*) crystals. MD (*orange*) and NMA (*purple*) mark the predictions from molecular dynamics simulations and normal mode analysis, respectively. Panels A and B show a comparison between the average properties of individual amino acid residues. Only approximately every 4^th^ residue is labelled to avoid crowding. Exceptions from this rule were allowed in order to highlight those residues that are mentioned in the discussion. Panels C and D compare different amino acid types.

Perhaps it is unrealistic to expect good agreement of U_ij_ derived values between experiments and predictions on an absolute scale. For example, the simulation was performed at 300 K instead of 100 K. Despite this inconsistency, B_eq_ was found to be on approximately the same scale. This can be explained by the fact that conformational heterogeneity is frozen in the crystal when it is flash-cooled to 100 K. Anisotropy, on the other hand, may be incorrectly estimated at much higher temperatures. Nevertheless, one expects that B_eq_ will be estimated self-consistently and reproducibly by MD simulations. Since Pearson correlation coefficients (CC) are not affected by the absolute magnitudes of compared values, they are also a suitable measure for comparing crystal structure and predicted values (self- and cross-correlations). As Table S2 shows, the MD B_eq_ estimations are not stable and their correlation with the estimated B_eq_ decays over time. It does not matter which reference time point is chosen. For example, the simulated trajectory from MD1 had a B_eq_ CC of 0.83, 0.71 and 0.64 when compared to MD2, 3 and 4, respectively. Choosing MD2 as a starting point also shows a decreasing CC trend. NMA B_eq_ shows a substantially lower correlation with any of the MD predictions (the best correlation is 0.45). Moreover, when compared to the B_eq_ of crystal structures, earlier trajectories show higher correlations than later ones irrespective of the crystal structure to which they are compared (Table S3). NMA B_eq_ correlations with crystals is on par with some of the MD predictions (the best correlation is 0.45 to crystal x28 and x30). Reference crystal structures had B_eq_ CC in the range of 0.99-0.82 among themselves due to experimental and modelling variation (Table S4).

Anisotropy values showed a smaller correlation within the experimental data and also between the theoretical predictions, although the predictions seem to become more stable over time (Tables S5, S6) than the CCs for B_eq_. Nevertheless, the combined experimental and modeling errors make predicted and experimental anisotropy hardly comparable (CC<0.4 for MD and CC<0.3 for NMA, Table S7). As such, we also have to consider the mismatch of the absolute level of anisotropy values between experiments and model predictions (Figure 4B). We provide additional discussion about the NMA and MD modelling in the Supporting Online Material.

### Metric-free clustering of atoms based on pairs of U_ij_ matrices

Hierarchical clustering was used in the crystallographic analysis, for example by grouping isomorphous crystals together (*32*) or detecting structural similarity.(*33*) In the context of analyzing ADPs, clustering algorithms have not been used before. Our atomic clustering was based on similarities between the components of ADP tensors, quantified by the Euclidean distance between the components (Figure 5). This way, atoms with similar shape, size and directionality are clustered together. The heat map represents the magnitude of the tensor components, providing quick visual feedback about the similarity and the nature of the remaining differences.

**Figure 5.**
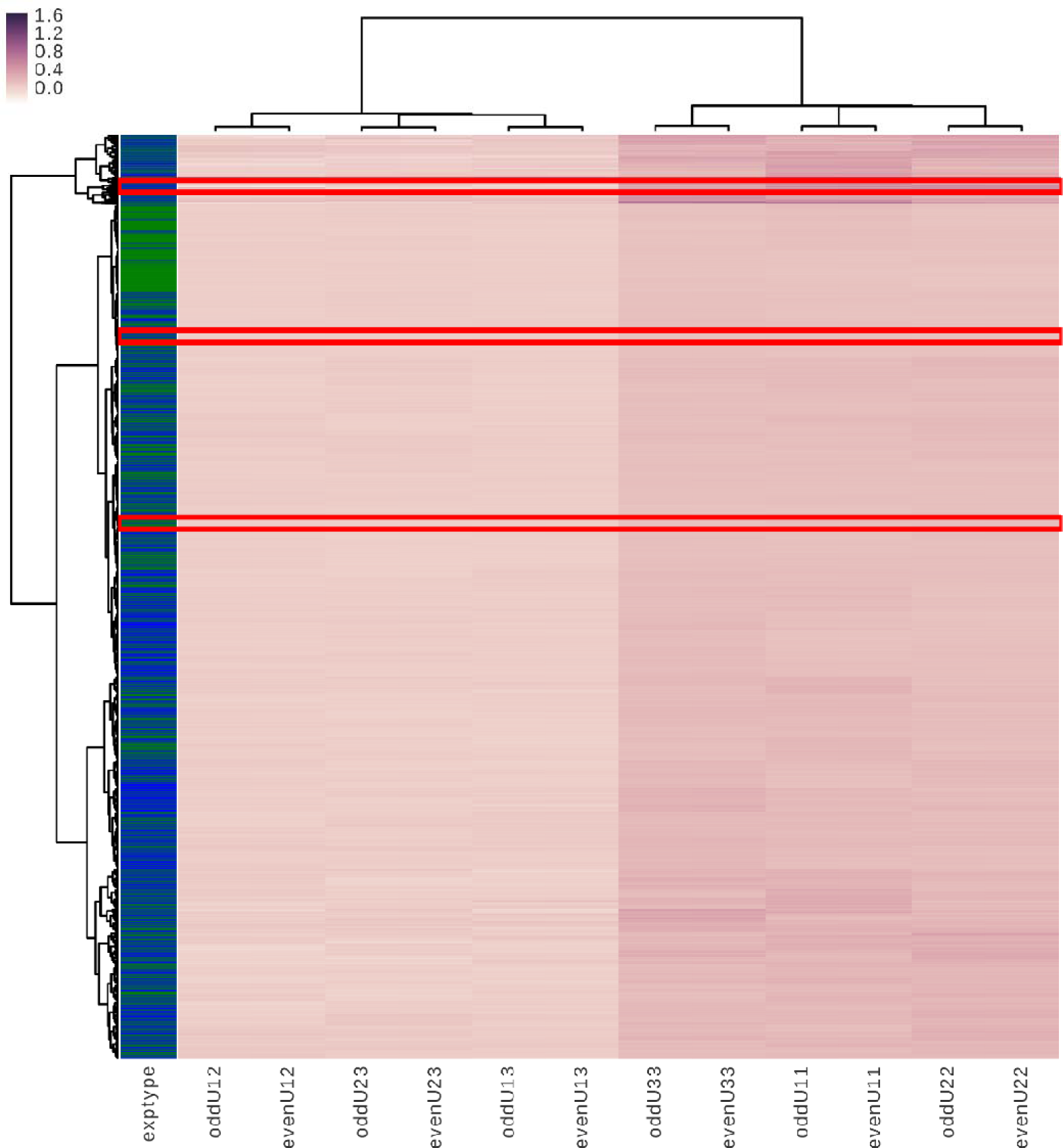
Cluster analysis of 15921 atoms of the two states (odd and even frames) of the four reference (*blue*) and five terahertz irradiated (*green*) crystals. Ideally, each line in the heat map and the corresponding green and blue color bar represent a single atom, but the resolution of this image is not sufficient to show all atoms. A more detailed view of the marked areas is shown in the subsequent plots. The dendrograms at the top and to the left represent the distances. The heat map represents the magnitude of each atom’s ADP tensor components (U_ij_) from the odd and the even state.

In Figure 5, the atoms from terahertz-radiated crystals are represented by green regions and these appear to cluster together. These clustered atoms originating from terahertz experiments tend to be located in the core regions of the protein structure. The atom mean displacement, represented by the sum of the diagonal elements (B_eq_), alone yields a very coarse clustering. However, slight differences in shape and orientation are enough to direct atoms to specific clusters.

In order to illustrate one type of formed clusters, Figure 6 zooms in on the top highlighted region in Figure 5. Here, we examine 7 out of 9 observations of the N_ε_ atom of Gln-135 on one branch. The immediately adjacent branch on the dendrogram consists of the N_δ_ atom of Asn-223, where all of the observations of this atom in all crystals can be found. It is worth emphasizing that no other information was used other than the 2 × 6 components of the U_ij_ matrices. Despite this, it is possible to uniquely identify up to 9 identical atoms from a total pool of 15921 atoms. Thus, ADPs are highly reproducible and weakly affected by experimental and modeling errors. It is important to note that these two atoms were not close to one other in the crystal structure, but they shared a similar position in a chemical group. The thermal ellipsoids these atoms are illustrated in Figure S2.

**Figure 6.**
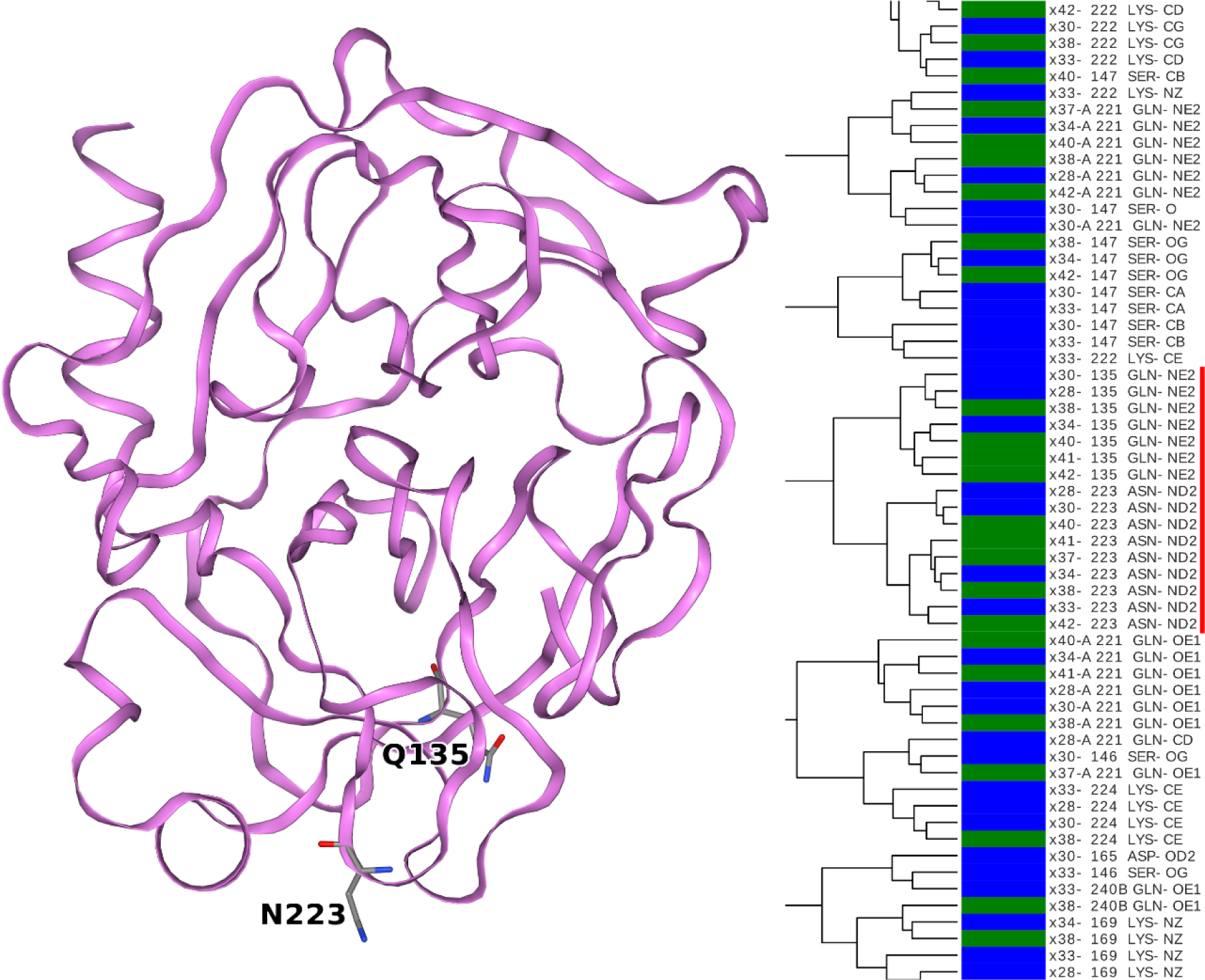
Clustering appears to identify amide nitrogen atoms of Gln-135 and Asn-223 residues in multiple crystals. (*red*) The two amino acids do not make intra- or intermolecular contacts. The trypsin structure is illustrated by a ribbon diagram. Those amino acid residues to which the clustered atoms belong are represented by sticks.

Figure 7 focuses on a central region highlighted in Figure 5, where most of the clustered atoms share the fact that they were observed in terahertz pumping experiments. The highlighted cluster contains three observations of C_β_ of Ser-195 and two observations of C_β_ of His-57. Ser-195 carries out the nucleophilic attacks and His-57 accepts the protons generated during the nucleophilic attacks. The ADPs of the two C_β_ atoms are illustrated as thermal ellipsoids in Figure S3. Atoms from Trp-215 and Ser-214 were also observed in this region. The backbone of these two amino acids form the antiparallel β-sheet with the substrate peptide in the Michaelis-complex and acyl-enzyme, and in addition the hydroxyl group of Ser-214 is hydrogen bonded to the third catalytic amino acid, Asp-102, which is not part of this cluster. The residue Trp-237 is represented with multiple atoms in the tree and this residue appears to be affected by terahertz radiation.

**Figure 7.**
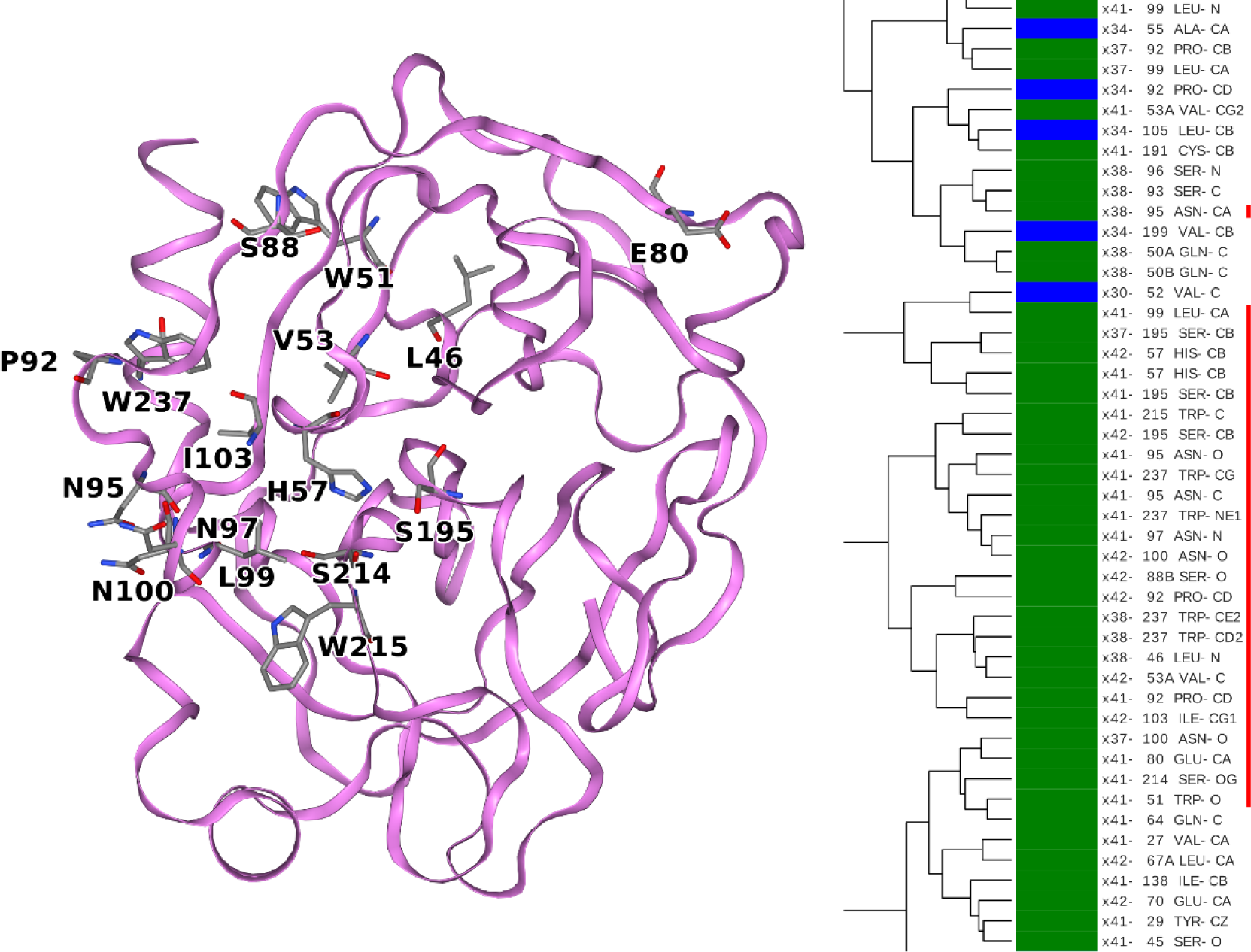
A cluster consisting of atoms from terahertz irradiated crystals that are involved in the catalytic process of the protein (Ser-195, His-57, Ser-214, Trp-215) and a high proportion of atoms originating from asparagine and tryptophan residues. The trypsin structure is illustrated by a ribbon diagram. Those amino acid residues to which the clustered atoms belong are represented by sticks.

Figure 8 shows the lower marked region in Figure 5, where the link between His-57 and Asp-102 appears to be stronger. In particular, there is a similarity between C_ε2_ of His-57 and O_δ2_ of Asp-102. In addition other O_δ2_ atoms of functionally important aspartates seem to be tuned together: Asp-194, which forms a salt bridge with N-terminal Ile-16 converting the enzyme into its active form, (*34*) and Asp-189, which (in general) forms the salt bridge with the P1 residue of the substrate. In this crystal structure, the Asp-189 forms a salt bridge with the amidinium group of the inhibitor benzamidine. It is important to mention that atoms belonging to Trp-237 and Trp-51 also appear in these trees, as the anisotropy of these tryptophanes is affected by terahertz radiation.

**Figure 8.**
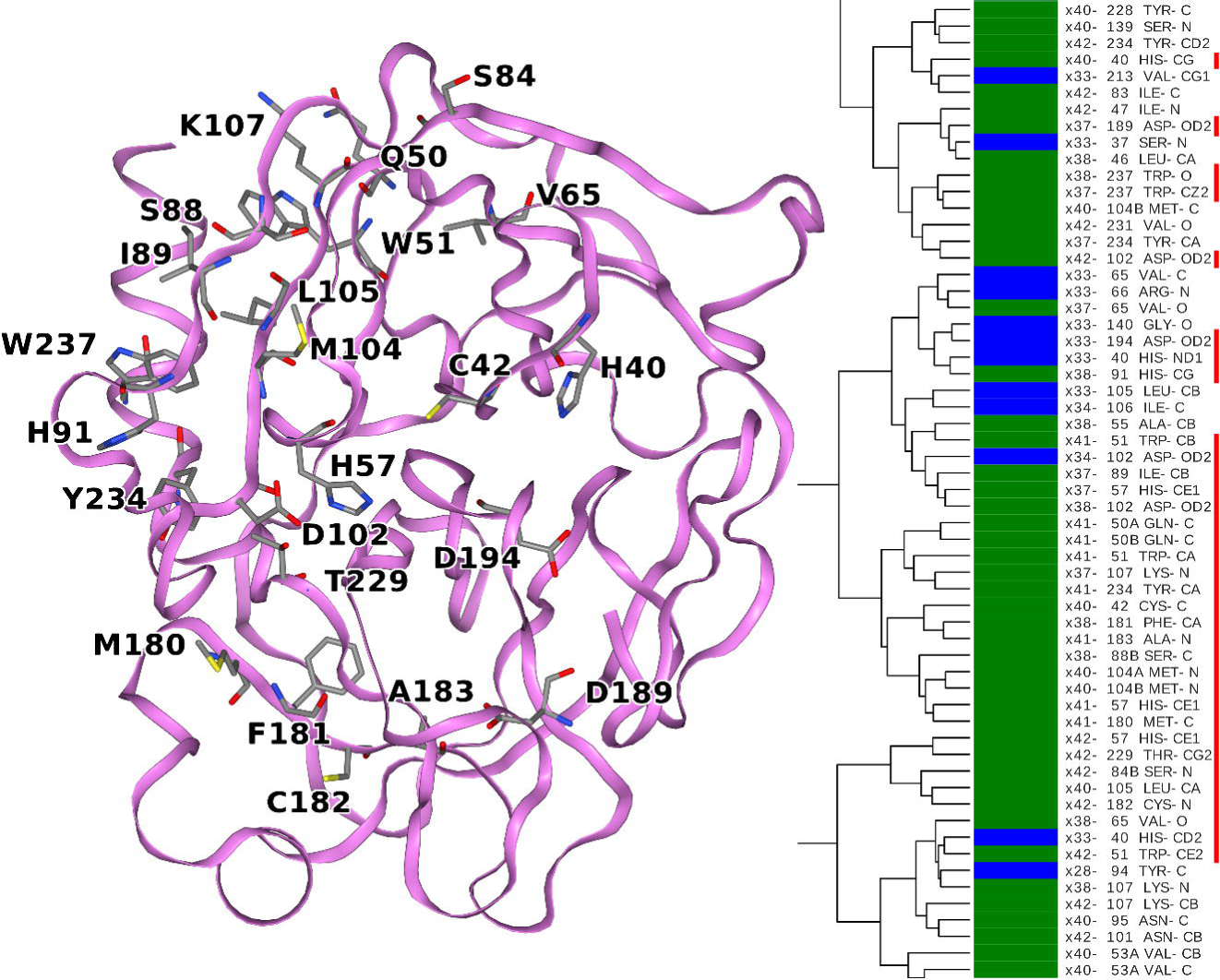
A cluster consisting of atoms from terahertz irradiated crystals that are involved in the catalysis and activation of trypsin (His-57, Asp-102, Asp-194 and Asp-189) a high proportion of atoms originating from tryptophan residues and all histidine residues are represented by atoms in the cluster. The trypsin structure is illustrated by a ribbon diagram. Those amino acid residues to which the clustered atoms belong are represented by sticks.

## Discussion

### The pattern of ADP clustering corresponds to trypsin function and reveals the dominance of polar vibrations

It is interesting to examine how the ADP clustering corresponds to the function of trypsin. Bovine trypsin is a serine protease that belongs to the chymotrypsin family. Its structure consists of two β-barrel domains that are stacked together and the catalytic triad of Ser-195, His-57 and Asp-102 amino acid residues are shared between them. (*35-37*) Trypsins cleave substrate peptides next to positively charged amino acid residues and the initial enzyme-substrate (Michaelis-) complex assumes an antiparallel β-sheet structure in which Ser-214, Trp-215 and Gly-216 participates from the enzyme’s side. (*38*) The first step is that the Ser-195 hydroxyl group commits to a nucleophilic attack on the carbonyl carbon of the scissile peptide forming a short-lived covalent tetrahedral intermediate. This intermediate then breaks down to the first product and the acyl-enzyme intermediate where the second product is tightly held by an ester bond to Ser-195. The second nucleophilic attack comes from an activated water molecule, which hydrolyzes the ester bond via a second tetrahedral intermediate. Both nucleophilic attacks are assisted by the His-57 - Asp-102 pair, which temporally accepts a proton from Ser-195, or from the activated water molecule, respectively. While the catalytic mechanism requires a subtle rearrangement of the active site atoms, there is a substantial energetic barrier to cross in order to overcome the remarkable stability of the peptide bond. The activation of the scissile peptide is partly provided by the polarizing effect of two main chain hydrogen bonds, termed the oxyanion hole, which is formed by the amide nitrogen atoms of Gly-193 and Ser-195. Longer substrates tend to be hydrolyzed faster which suggests that collective motions of the substrate-enzyme β-sheet contribute to the catalytic efficiency. (*38*) On the surface, it appears that the mechanism of trypsin catalysis is well understood, but we still lack a predictive model that associates a mutation or allosteric modulation with an expected change in catalytic parameters and substrate specificity. (*39*)

Atoms in functionally important residues appear repeatedly together in local clusters. Atoms in separate branches are often predominantly observed in either terahertz radiated or non-radiated crystals, respectively. In Figure 7, one such terahertz radiated branch (green) is highlighted. A large physical distance between these atoms does not appear to be a hindrance, whereas the chemical nature of the atoms clearly does have an influence. For example, not all carbon atoms are preferentially associated with other carbon atoms, but carbon atoms in carbonyl, α and β positions in the amino acid residues are different enough to sort them to different clusters. (Figure S4) One could consider the ADP similarity to be an effect of the atom types having a similar partial charge. This could be a tempting assumption for the amide side chain nitrogen of asparagine and glutamine (Figure 6), which are assumed to have a strong and similar negative partial charge in most empirical force fields. It would also make sense that atoms with a similar partial charge respond similarly to a uniform electrostatic field. In a sodium chloride crystal, there are two well-defined charged ions (Na^+^ and Cl^−^), and these respond to electromagnetic (EM) fields in two possible ways. As a result, polar vibrations (optical phonons) emerge. In a protein, there is a whole palette of atom types, and we approximate each as having a specific static partial charge between approximately −0.8 and +0.9. Consequently, each of these atom types may have a slightly different response to EM fields. On the other hand, the origin of the terahertz EM field makes no difference. The EM field may be generated internally by the protein (crystal), by the solvent, by external pumping or any combination of these. Since the wavelength of terahertz EM waves is much longer than the size of protein molecules they generate virtually uniform EM fields across the protein molecule at any given moment. On the other hand, the partial charges of C_β_ atoms are not necessarily similar. For example, C_β_ of Ser-195 is assumed to be partially positive because of its bonding to the more electronegative hydroxyl group, whereas C_β_ of His-57 is assumed to be more neutral, because it is attached to another carbon of an imidazole ring. Nonetheless, these two C_β_ atoms reproducibly cluster together (Figure 7).

### Do protein domain structures emerge from entrainment of atomic displacements?

In architecture, one models the dynamics of an arbitrary structure (a house or a bridge) and then alters the structure until the dynamical properties are acceptable. Molecular NMA follows this same conventional logic, except that making planned changes in a protein structure is not a trivial process. In contrast to conventional engineering, a protein structure is not created by placing the atoms into arbitrary positions; rather, it emerges spontaneously from a quite dynamic polymer under suitable conditions (protein folding). The ADPs of atoms do not (exclusively) depend on where they are located in the structure, which suggests that most atoms are not passive passengers in their structural elements.

We show evidence here that in the folded, crystallized form of a protein, atomic fluctuations are characteristic, stationary (reproducibly seen in multiple crystal structures) and coordinated across long distances connecting together different structural features. At first glance, intriguingly, atoms of a sharply defined chemical nature align themselves with similar atoms in other identical chemical groups. Another non-random aspect is that atoms cluster together when they originate from terahertz pumped crystals (Figure 5 and 7). The dissipation of applied terahertz radiation enhances the clustering of atoms for example by grouping identical atoms from multiple crystals into the same cluster and placing atoms from functional residues to common groups. On the other hand, the ADPs in our reference crystals already provide substantial information about the long-range pattern: it is not exclusively present in an out-of-equilibrium condition.

In a protein crystal structure some, but not all, chemically similar atoms have similar ADPs. The mechanism for selecting from the many chemically similar atoms needs to be investigated further. It appears that protein atoms form an intercalating mosaic of coexisting lattices, where it is common to see atoms from the same amino acid residues ending up in different clusters. Perhaps the most noticeable, but certainly not exclusive, example is the group of tryptophan residues, where the indole ring is often approximated as a rigid aromatic plane with torsional fluctuations around two axes as the most important motions. Complicating this picture, the atoms within the indole ring cluster differently and share clusters with atoms in other tryptophan residues. Tryptophan residues are often difficult to replace by mutagenesis and play a key role in maintaining the thermodynamic stability of protein structures. They are one of the strongest promoter of ordered state. (*40, 41*) They also appear to be reversibly affected by terahertz radiation in this study.

The disorder-to-order transition is also facilitated by asparagine and threonine residues. (*41*) In particular asparagine residues were poorly described by MD simulations, which predict larger B_eq_ and lower ANISO in absolute terms and relative to other amino acid types (Figure 4C and 4B). It appears that in the folded form, the side chain amides of asparagine residues undergo very limited fluctuations even in the absence of local interactions. Given the limited local contacts, MD simulations predict more positional freedom. In the terahertz experiments, some of the asparagine atoms also appear to clusters together with catalytic residues (Figure 7). This is probably more the exception than the rule, since asparagines and glutamines in general do not appear to be significantly affected by external pumping (Figure 2, Figure 4C and 4B). Asparagines do not readily maintain a periodic secondary structure in contrast to their chemical relative, glutamines. (*42*) Glutamine side chains have greater orientational freedom with multiple potential alignment possibilities and can presumably support multiple alternative stationary conformations leading to an increase in long-range disorder. A preliminary proposal is presented on how ADP similarity might be linked to structural constraints and to the symmetry between chemical groups in a protein in the Supporting Online Discussion, Figure S4 and Figure S5.

Evolutionary processes change the chemical composition of proteins and thus change protein structure very slowly. In contrast, post-translational and epigenetic modifications can introduce multiple chemical groups of identical nature (phosphorylation, sulfurylation, acetylation, methylation, etc.) rapidly. Presumably, these post-translational modifications interact in a similar manner and redefine the (stationary or non-stationary) positional distributions of atoms depending on the pattern of the modifications. A particular example is the nucleosome histone subunits (H3, H4), each possessing highly dynamic N-terminal regions that are subject to epigenetic regulation through a combination of phosphorylation, methylation and acetylation at specific amino acid residues. Each combination of modifications has a finely tuned functional consequence, which several nucleosomes carry out in a concerted manner with considerable distance between them. These modifications include promotion and silencing of gene transcription, unpacking of DNA and the formation of heterochromatin.

### Delocalized polar vibrations are important for structure and function prediction

Physical proximity and local complementarity have been and still are the primary guiding principles in protein engineering. Local shape complementarity is necessary, but not sufficient condition for structure formation. Directed engineering of protein interfaces is less than straightforward due to the elusive (allosteric) influence of the rest of the structure. The similarity of parameters that describe fluctuations are not routinely analyzed, however. B_iso_-factors and NMR relaxation parameters by themselves do not contain much information: two atoms can easily have similar parameters, just by coincidence. Experimental and modeling errors make the comparison even more difficult. It requires series of observations (*43, 44*) in order to recognize the covariance of these parameters and assign multiple atoms to a specific functional role. With anisotropic ADPs, the chances of coincidental similarities are dramatically reduced. While multiple related structures help (as demonstrated in this study), it is possible to detect similarities between atoms with good confidence even in a single crystal structure. The examples described here show that the anisotropy of these fluctuations appears to have a functional relevance with two or more atoms with similar ADPs being responsible for a similar function (catalysis, conformational change between active and inactive form of the enzyme). These similarities can be revealed faster than time-consuming mutagenesis studies, which are also impossible when main chain atoms are concerned. While the extracellular enzyme trypsin is not typically seen as an allosteric enzyme, its substrate specificity, activation, and inhibitor binding involve diffusely distributed, concerted action of distant amino acid atom positions suggesting that the basis of allostery is a universal, emergent and dynamic process.

Biomolecular disorder incorporates diffusive motions of folded and unfolded domains relative to one other (Figure S6), conformational changes, elastic stretching (Figure S6A), bending (Figure S6C) and torsions. It appears that that a rigid body approximation, even at the level of small structural moieties such as amino acid side chains, does not provide natural clusters for ADP similarities. Our study demonstrates that within a folded unit polar terahertz vibrations can be identified (Figure S6D). Importantly, these delocalized vibrations may have the ability to steer the orientation of involved atoms at great distances and the electron dynamics of covalent bonds provide additional local constraints for orienting the attached chemical moieties. The identification of similar ADPs in a macromolecular structure is just the first essential step. The second and possibly more difficult step is to find and understand the geometric patterns between atoms belonging to the same ADP cluster. Since similar ADPs seem to be linked to the chemical nature of atoms, empirical geometric rules may be recognized or applied in the absence of ADP observations. These patterns would then assist the development of long-range ADP restraints, complementing restraints based on bonding connectivity and positional proximity. Improved ADP restraints may provide better electron density maps and more detailed structural models at lower crystallographic resolutions. More importantly, long-range geometric patterns between chemically similar atoms coupled to effective conformational sampling (simulated annealing) may be able to assist the *de novo* prediction of protein structures.

## Conclusions

Our study focused on two inseparable questions: the effect of terahertz radiation on protein crystals and the terahertz dynamics of folded proteins. We used crystallographic ADPs to measure the effect of terahertz radiation on crystals cooled to 100 K. Locally, we observed an increase in anisotropy and we focused our attention on glycine and tryptophan residues, and the catalytic residue His-57, where the most obvious changes in anisotropy occur. The clustering of atoms based on their ADPs provided a relational map, which complements the positional analysis of atoms based on their proximity. We performed state of the art NMA and MD simulations to predict ADPs and achieved limited success only. The clustering revealed that chemically similar atoms tend to have ADPs that are more similar. These atoms are distributed sparsely in the protein structure and belong to functionally connected amino acid residues.

## Methods

### Protein crystallization and flash cooling

Bovine trypsin (Sigma) was dissolved in 30 mM HEPES pH 7.0, 3 mM CaCl_2_ and 6 mg/ml benzamidine to obtain 60 mg/ml protein solution. Trypsin was crystallized using the hanging drop vapor diffusion method, by mixing 5 μl protein solution and 5 μl of precipitant solution (18% PEG8000, 50 mM HEPES pH 7.0, 0.2 M Ammonium sulfate, 3 mM CaCl_2_ and 6 mg/ml benzamidine). After 3 days, orthorhombic crystals appeared with approximate dimensions of 100-300 µm. The crystals were harvested and soaked in cryoprotectant solution (additional 33% glycerol) for 1-2 seconds prior to flash cooling in liquid nitrogen.

### X-ray diffraction data collection

X-ray diffraction data were collected at the EMBL P14 beamline in Hamburg. The incident X-ray beam was focused to a 10 µm horizontal × 50 µm vertical rectangular area at the sample position. The photon energy of the beam was 14 keV and the beam was attenuated to a photon flux of 4 × 10^11^ photons/s. The central panels of the Pilatus 6M-F detector were used (2463 × 2527 pixels) and the detector was placed at the distance of 130.4 mm from the crystal. We discuss the effect of X-ray radiation damage on ADPs in the Supporting Online Material and Figure S1.

The crystals were oriented with the minikappa goniometer such that the normal vector of clean crystal surface was aligned with the spindle direction. The beam cross section was positioned less than 100 μm from the crystal surface and 20 μm offset from the spindle axis. During X-ray exposure the crystals were continuously rotated with 0.4 °/s while the Pilatus 6M-F was continuously recording images at 25 ms intervals. During each image recording period, the detector spent 22 ms with photon counting and 3 ms with I/O operations (read out).

The terahertz radiation was generated by an amplifier / frequency multiplier chain (X32 stage AMC, manufactured by Viginia Diodes Inc., Charlottesville, VA), which was driven by a MG3692c microwave signal generator (Anritsu). The input microwave radiation was set to 15.625 GHz (10 mW), which resulted in 0.5 THz radiation (1 mW). The diagonal horn antenna (WR-2.2) was positioned approximately at 0.5 cm from the crystals. Under the assumption of a Gaussian beam profile, the THz spot size was approximately 4 mm. This estimate assumes a waist radius of 1.3 mm and that the beam originates from approximately 1/3 inside the antenna. We examine the thermal effect of the terahertz radiation in the Supporting Online Material.

To reduce systematic differences associated with thermal effects, the terahertz radiation source was operated alternately. A DG645 pulse (delay) generator (Stanford Research Systems) was triggered on the raising edge of the EN OUT signal of the Pilatus 6M-F detector. Upon triggering, the pulse generator was programmed to send a 23.5 ms pulse to the terahertz radiation source (THz_on_ period). Simultaneously, the pulse generator was set to ignore subsequent triggers for 40 ms. Consequently, every odd numbered image was taken from the crystal in the terahertz irradiated state whereas the even numbered images recorded the diffraction image of the crystal in the non-irradiated state.

Odd and even numbered diffraction images still contained sufficiently super sampled measurements to integrate their total intensity independently. (*11, 45*) This applies even for the widest-angle reflections recorded for this study, observed far away from the spindle axis.

### X-ray diffraction data analysis

The first diffraction image was always affected by the slow opening of the millisecond shutter and this image had systematically lower recorded scattering intensity. The data processing program XDS (*46*) uses the first image of each data set for initial scale, which introduced a systematic difference between the data sets based on odd and even images. Therefore, the first odd and first even images of each data set were removed from subsequent analysis. In addition, the kappa goniometer shadow was identified in each data set and the affected images were removed. Even number of images were removed in order to maintain the equal number of odd and even numbered diffraction images.

The sets of diffraction images, from the odd and even frames of each crystal were processed separately. By comparing the data and refinement statistics, we selected the crystals with generally high data quality, shown in Table S1. This pre-selection yielded 18 data sets, recorded from four reference and five terahertz radiated crystals. Moreover, since the frames affected by the goniometer shadow differed from crystal to crystal, the unaffected rotation wedges were integrated individually. The indexing and integration steps were carried out with the default settings in XDS. The subsequent scaling and merging were performed using the programs XSCALE and XDSCONV of the XDS package, respectively. In both programs, the standard settings for non-anomalous diffraction data were used.

Similarly, the subsequent model building and structural refinement were performed on each individual state (the odd and the even frames) of the individual crystals, using Refmac5 of the CCP4 package. (*47*) A previously refined model was used as a basis for the Refmac5 refinement (see Supporting Online Information).

### Model building and structural refinement

A single starting model was refined against the 18 data sets (9 crystals, recorded on odd and even diffraction images). Refinement statistics is shown in Table S1.

The initial structure was based on pdb entry 4I8G, and the structure was solved by molecular replacement. (*48*) The starting model included 2058 non-hydrogen atoms 111 of which belonged to water molecules, 15 were heteroatoms and 1932 were part of amino acid residues. Riding hydrogen atoms were included during refinement. The initial refinement leading to a starting model is described in the Supporting Online Information.

The individual crystal refinement was carried out with Refmac5, using the rigid body, restrained isotropic and restrained anisotropic refinement procedures. Each refinement procedure was performed with the default settings for 100 cycles with an automatic weighting of restraints.

### ADP analysis and clustering

The calculations and visualization were performed with the help of the python libraries numpy, scipy, pandas,(*49*) cctbx,(*50*) seaborn and NGLView. (*51*) Through hierarchical clustering, all the observed atoms were iteratively grouped using the distance between their unmodified U_ij_ matrices. By having two observations of each atom in the same crystal, we obtained two symmetric U_ij_ matrices with six unique elements each. The clustering was thus performed in a twelve-dimensional data space and we used the Euclidian distance between these points. For clustering, Ward’s method was used. (*52*)

### Apparent X-ray dose calculations

In order to compare the apparent absorbed doses in the different crystal structures first we estimated the cumulative absorbed X-ray dose corresponding to each diffraction image. The absorbed dose was calculated using the program RADDOSE 3D. (*53*) The apparent dose was calculated as the product of the dose per frame, multiplied with the weighted arithmetic mean of the number of frames for each crystal.

### Normal mode analysis and molecular dynamics simulations

Normal mode analysis (NMA) was performed using the Gromacs simulation package (*54-56*) version 2018-2, compiled with double precision. The structure was first energy minimized rigorously by employing several cycles of Steepest Descents and Conjugate Gradients energy minimization cycles. The CHARMM36 (*57-59*) force field was used in all calculations. Electrostatics and van der Waals forces were truncated at 1.8 nm with a shift cutoff from 1.5 nm. After energy minimization, the Hessian matrix was diagonalized and all eigenvalues and eigenvectors were used except the first six rotational and translational modes. An ensemble of structures was generated at 100 K by the Gromacs program gmx nmens and ADPs were calculated by the Gromacs program gmx rmsf for all non-hydrogen protein atoms.

Molecular dynamics simulations were carried out using energies and forces as implemented in the OPLS-AA/L force field. (*60*) The protein was solvated in a cubic box so that the minimum distance between any protein atom and the edge of the box was 1.0 nm. The water molecules were modeled with the TIP3P approach. (*61*) Cl^−^ ions were added in order to neutralize the charge. Before the MD simulation, internal constraints were relaxed by energy minimization. After the minimization, a restrained MD run was performed for 20 ps. During the restrained simulations, protein heavy atoms were fixed to their initial positions with a force constant of 1000 kJ mol^−1^ nm^−2^. The restraints were released and the system was equilibrated for 60 ns before data collection for analysis.

The temperature was kept constant (T=300K) by use of the velocity-rescaling algorithm (τ_T_=0.1 ps). (*62*) The pressure was coupled to an external bath with Berendsen’s coupling algorithm(*63*) (P_ref_=1 bar, τ_P_=1 ps) during the equilibration and with the Parrinello–Rahman algorithm(*64*) afterwards. Van der Waals forces were truncated at 1.0 nm with a plain cutoff. Long-range electrostatic forces were treated using the particle mesh Ewald method.(*65*) Dispersion correction was applied for the energy and pressure.

After equilibration, the remaining 40 ns of simulations were divided into 10 ns segments and the ADPs were calculated with the Gromacs program gmx rmsf from the trajectory segments.

## Acknowledgements

This work was supported by the Swedish Research Council, the Röntgen-Ångström Framework and the Knut and Alice Wallenberg Foundation. The diffraction experiments were performed at the P14 beamline of Petra III.

