## Supporting Online Material for "A distributed lattice of aligned atoms exists in a protein structure: A hierarchical clustering study of displacement parameters in bovine trypsin"

Viktor Ahlberg Gagnér^1^, Ida Lundholm^1^, Maria-Jose Garcia-Bonete^1^, Helena Rodilla^2^, Ran Friedman^3^, Vitali Zhaunerchyk^4^, Gleb Bourenkov^5^, Thomas Schneider^5^, Jan Stake^2^ and Gergely Katona^1,*^
^1^ Department of Chemistry and Molecular Biology, University of Gothenburg, Gothenburg, Sweden.

^2^ Department of Microtechnology and Nanoscience, Chalmers University of Technology, Gothenburg, Sweden.

^3^ Department of Chemistry and Biomedical Sciences, Linnaeus University, Kalmar, Sweden

^4^ Department of Physics, University of Gothenburg, Gothenburg, Sweden.

^5^ European Molecular Biology Laboratory Hamburg Outstation, EMBL c/o DESY, Notkestrasse 85, 22603 Hamburg, Germany.

Supporting Online Results

*The effect of X-ray radiation damage on B_eq_ and ANISO metrics*

Although the crystals received approximately equal X-ray doses, collecting high-resolution (wide-angle) diffraction data can be associated with more frequent incidents of shadows cast by the minikappa goniometer. These moving shadows can interfere with the diffraction intensity estimation; therefore, the affected images were removed. Since shadows appear at different rotation angles during each experiment, the excluded diffraction images remove information about the associated radiation damage state of the crystal. When merging the remaining intensity observations, the Bragg reflections display an apparent X-ray radiation damage that is different from the total or average dose deposited on the crystal. Thus, we base the estimate of the observed radiation damage level on the apparent absorbed X-ray radiation dose of the retained images of each crystal.

The absorbed dose was estimated using the RADDOSE-3D web interface. As a first approximation, we estimated the upper limit of absorbed X-ray dose by assuming that the crystal is stationary for the same time duration as one data collection (360° rotation). This yielded 9.7 MGy as the total dose spatially averaged over the exposed region. Based on this limit, the reference set of crystals reported 2.7 ± 0.30 MGy as the average apparent dose. The ± represents the standard error of the mean. For the terahertz irradiated data, the average apparent dose were 2.4 ± 0.30 MGy. A more realistic estimate of the total dose, given by RADDOSE during a rotation of 360°, was 1.3 MGy. Thus, the reference and terahertz irradiated data sets show an average apparent radiation damage corresponding to 0.36 ± 0.04 MGy and 0.32 ± 0.04 MGy, respectively.

Additionally, we used RADDOSE-3D to calculate the total amount of dose per frame, to compare the difference in absorbed dose between the terahertz irradiated and non-irradiated data sets (the odd versus the even frames). The doses per frame were calculated as 270 Gy/frame and 40 Gy/frame for the pessimistic and the realistic case, respectively. Considering the fact that the data collection starts with and odd and ends at an even frame, the even data set receives a radiation damage attributed to 40-270 Gy more, compared to the odd set.

The general effect of X-ray radiation damage to protein crystals is the ionization of atoms and the breaking of atomic bonds. For sufficient absorbed X-ray doses, the effect of radiation damage is noticeable in for instance crystal model parameters, such as an increase in the crystal volume and in atomic B-factors. (*1*) The apparent dose difference between the reference and THz irradiated data sets are calculated to be 0.30 ± 0.85 MGy, and 0.04 ± 0.11 MGy, in the pessimistic and realistic case, respectively. Since the apparent X-ray dose difference between the reference and terahertz irradiated crystals are not substantial in comparison to the Henderson limit (20 MGy),(*2*) the differences in the model parameters are not likely related to the difference in apparent absorbed dose. This especially applies also to the dose difference of the odd versus the even data sets. Additionally, Shimizu et al. (*3*) show that the magnitudes of the calculated doses does not substantially affect the protein structure. The authors show that the change in B-factors could be linear for small X-ray doses. Therefore, we assumed this relationship for change in our B_eq_-factors. From the resulting plot (Figure S1), we obtained a change corresponding to approximately 3.07 Å^2^ / MGy from the reference data set. By considering the differences in absorbed apparent doses, the estimated change in B_eq_ due to X-ray radiation damage is 0.92 ± 2.61 Å^2^ in the pessimistic case, and 0.12 ± 0.34 Å^2^ in the more realistic case. Since the average difference of the B_eq_ from the odd state of the terahertz and reference data sets is approximately 1.08 ± 0.56 Å^2^, the difference is not likely only related to X-ray radiation damage. When considering the general expression for atomic ADP’s, there seem to be a general trend that they become more anisotropic. Certain tensor parameters increase at a different rate. (*4, 5*)

Additionally, a larger variance in the average B_eq_-factors is shown in Figure S1 for the terahertz irradiated crystals, which results in a generally lower Goodness of Fit. The larger variance indicates that an external source could perturb the B-factors.

[1]. *Estimation of the temperature change induced by the terahertz source*

Upon terahertz irradiation, the temperature increases (up to a steady state). As a result, the magnitudes of the ADP’s may increase as well. ADPs are also called temperature factors in crystallography, because displacement may be the result of temperature-dependent atomic vibrations. The observed decrease in B_eq_ is not expected as a thermal effect since a temperature increase is almost exclusively associated with increased crystal B-factors.(*6*)

The temperature change due to terahertz irradiation can be modeled as follows: by considering the Gaussian part of the radiation and the antenna specifications, the power density of the wave front 5 mm away from the antenna (distance used in the experiment) was around 6.5 mW/cm^2^. The trypsin crystal was approximated as ice. During the experiment, the crystals were under the flow of a cryostream at 100K. This is translated into a permittivity with real part 3 and imaginary part of 0.005 at 0.5 THz. (*7*)

In the calculation it is assumed that the terahertz beam impacts perpendicularly to a flat surface of a hypothetical crystal. The crystal face had an area of 9×10^4^ µm^2^ and the absorbed power in the first 50 µm of the crystal was 8 nW (reflections due to change of media have been considered). The absorbed energy was equivalent to approximately one 0.5 THz photon per two second per unit cell. Ice at 100 K has specific heat capacity of 3.8 cal/mol/degree. (*8*) During the 23.5 ms that the terahertz source is “on”, the estimated absorbed power gives rise to a temperature increase of around 54 µK. This is the estimated temperature fluctuation between the constant minimum and maximum temperature in a duty cycle (steady state assumption). The minimum and maximum steady-state temperatures are expected to be reached in 1-3 seconds. During the steady-state phase of the experiment, the adjacent images showed a crystal with very similar average temperature (the “on” states correspond to an increasing temperature and vice versa for the “off” states). During the detector read out period, diffraction was not observed (3 ms), therefore the high and low temperature state of the crystal was incompletely recorded. However, since the terahertz pulse finishes in the middle of the read-out period, the unobserved period started and finished at approximately the same high temperature in the steady-state phase of the experiment. It is important to note that the steady state assumption implies that the heating and cooling rates are slightly different when the duty cycle of the terahertz source is 47%.

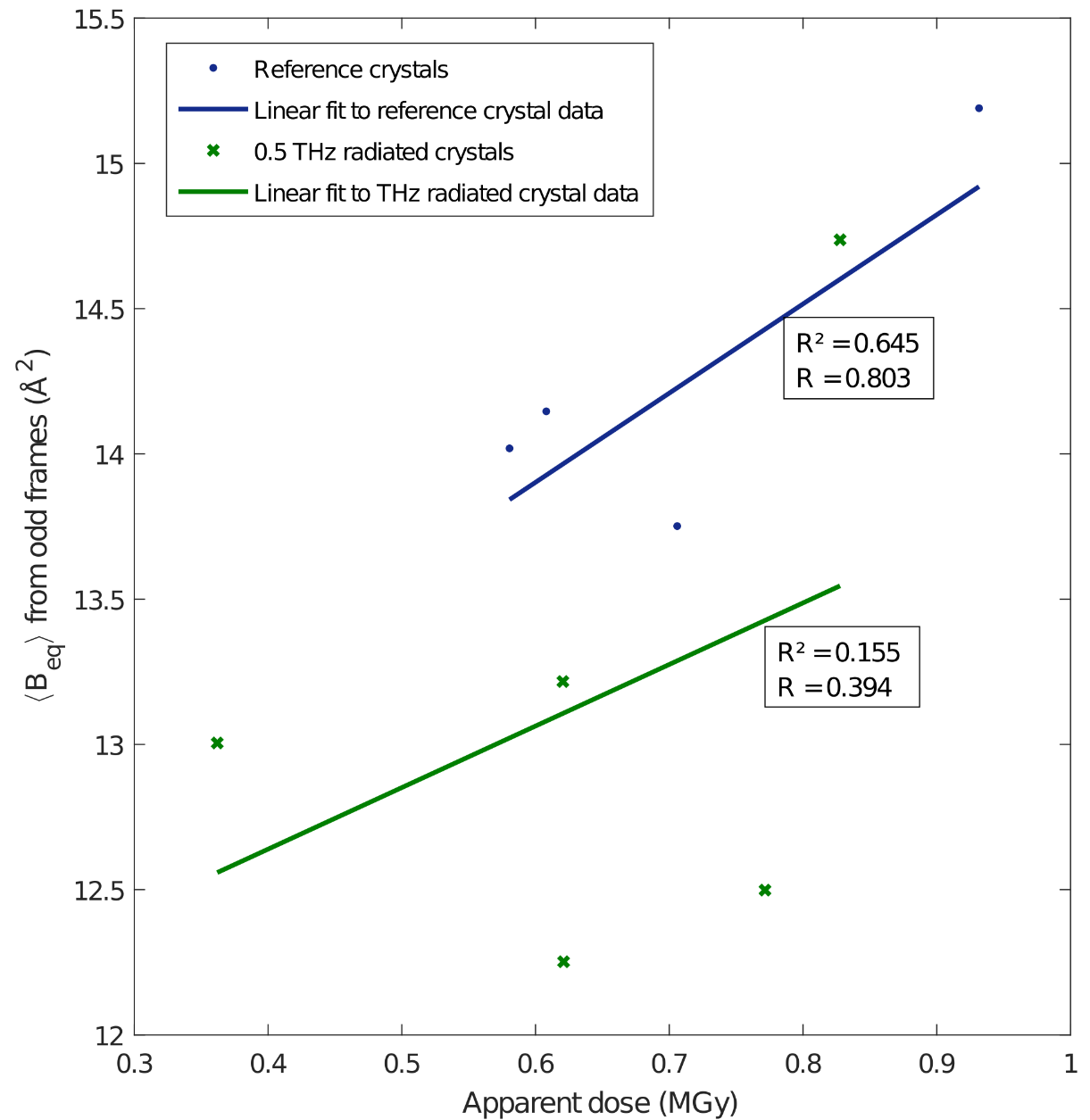

**Figure S1** Average B_eq_ from odd frames in the reference (*blue*) and terahertz radiated crystals (*green*), plotted versus the apparent absorbed X-ray doses.

**Table S1** X-ray data collection and refinement statistics. The values are obtained as means from four reference crystals and five terahertz radiated crystals after the scaling step. The mean B-factor refer to the average B-factor (B_eq_) of all atoms in the crystals.(*9*)

|  | Odd frames reference crystal  (no THz) | | Even frames reference crystals (no THz) | Odd frames  0.5 THz crystals (THz on) | Even frames  0.5 THz crystals (THz off) |
| --- | --- | --- | --- | --- | --- |
| **Data collection:** | |  | | | |
| Space group: | | P 2_1_2_1_2_1_ | | | |
| Cell dimensions  a; b; c (Å):  α; β; ɣ (°): | | 53.39 ± 0.04  56.78 ± 0.03  65.06 ± 0.11    90;90;90 | 53.39 ± 0.04  56.78 ± 0.03  65.06 ± 0.11    90;90;90 | 53.53 ± 0.11  56.90 ± 0.11  65.48 ± 0.13    90;90;90 | 53.53 ± 0.11  56.90 ± 0.11  65.48 ± 0.13    90;90;90 |
| Resolution (Å): | | 42.82 - 1.15 | 42.82 - 1.15 | 43.03 - 1.15 | 42.97 - 1.15 |
| I / σ*: | | 10.6 ± 1.8  (1.0 ± 0.2) | 10.6 ± 1.8  (1.0 ± 0.2) | 8.8 ± 0.63  (1.0 ± 0.1) | 8.8 ± 0.6  (1.0 ± 0.1) |
| Completeness (%)*: | | 94.0 ± 1.0  (41.8 ± 4.5) | 94.0 ± 1.0  (41.6 ± 4.6) | 95.0 ± 0.4  (46.6 ± 2.5) | 95.0 ± 0.4  (46.4 ± 2.5) |
| Redundancy*: | | 4.5 ± 0.4  (2.0 ± 0.2) | 4.5 ± 0.4  (2.0 ± 0.2) | 4.3 ± 0.4  (1.8 ± 0.13) | 4.3 ± 0.4  (1.8 ± 0.13) |
| CC_1/2_*: | | 99.8 ± 0.1  (44.0 ± 8.4) | 99.8 ± 0.1  (41.7 ± 9.1) | 99.7 ± 0.1  (37.8 ± 10.9) | 99.7 ± 0.1  (37.3 ± 10.6) |
| R_work_ / R_free_ (%)*: | | 13.4 ± 0.2 /  17.8 ± 0.6  (30.0 ± 2.8 /  34.0 ± 2.6 ) | 13.7 ± 0.4 /  18.0 ± 0.8  (30.0 ± 3.0 /  33.0 ± 2.8) | 13.7 ± 0.4 /  17.7 ± 0.5  (32.2 ± 2.4 /  33.8 ± 2.2) | 13.7 ± 0.4 /  18.0 ± 0.5  (32.3 ± 2.3 /  34.9 ± 1.8) |
| Mean B-factor (Å^2^): | | 14.23 ± 0.31 | 14.32 ± 0.33 | 13.15 ± 0.43 | 13.13 ± 0.43 |

*The values in the parentheses are taken from the highest resolution bin (from 1.18-1.15 Å).

**Table S2** Pearson correlation coefficients between per residue averages of B_eq_ values. Symmetrical pairwise comparisons were performed between four parts of an MD trajectory and NMA estimate.

| **CC B_eq_** | **MD 1** | **MD 2** | **MD 3** | **MD 4** | **NMA** |
| --- | --- | --- | --- | --- | --- |
| **MD 1** | 1.00 | 0.83 | 0.71 | 0.64 | 0.42 |
| **MD 2** | 0.83 | 1.00 | 0.68 | 0.58 | 0.40 |
| **MD 3** | 0.71 | 0.68 | 1.00 | 0.74 | 0.45 |
| **MD 4** | 0.64 | 0.58 | 0.74 | 1.00 | 0.38 |
| **NMA** | 0.42 | 0.40 | 0.45 | 0.38 | 1.00 |

**Table S3** Pearson correlation coefficients between per residue averages of B_eq_ values. Pairwise comparison was performed between four parts of an MD trajectory, the NMA estimate and the experimentally determined B_eq_ values of the reference crystals (odd frames).

| **CC B_eq_** | **x28** | **x30** | **x33** | **x34** |
| --- | --- | --- | --- | --- |
| **MD 1** | 0.61 | 0.60 | 0.60 | 0.56 |
| **MD 2** | 0.49 | 0.48 | 0.44 | 0.45 |
| **MD 3** | 0.46 | 0.44 | 0.39 | 0.44 |
| **MD 4** | 0.38 | 0.37 | 0.33 | 0.37 |
| **NMA 5** | 0.45 | 0.45 | 0.41 | 0.41 |

**Table S4** Pearson correlation coefficients between per residue averages of B_eq_ values. Symmetrical pairwise comparisons were performed between four reference crystals (odd frames).

| **CC B_eq_** | **x28** | **x30** | **x33** | **x34** |
| --- | --- | --- | --- | --- |
| **x28** | 1.00 | 0.99 | 0.93 | 0.96 |
| **x30** | 0.99 | 1.00 | 0.95 | 0.93 |
| **x33** | 0.93 | 0.95 | 1.00 | 0.82 |
| **x34** | 0.96 | 0.93 | 0.82 | 1.00 |

**Table S5** Pearson correlation coefficients between per residue averages of ANISO values. Symmetrical pairwise comparisons were performed between four parts of an MD trajectory and NMA estimate.

| **CC ANISO** | **MD 1** | **MD 2** | **MD 3** | **MD 4** | **NMA** |
| --- | --- | --- | --- | --- | --- |
| **MD 1** | 1.00 | 0.83 | 0.69 | 0.69 | 0.31 |
| **MD 2** | 0.83 | 1.00 | 0.70 | 0.65 | 0.34 |
| **MD 3** | 0.69 | 0.70 | 1.00 | 0.76 | 0.35 |
| **MD 4** | 0.69 | 0.65 | 0.76 | 1.00 | 0.37 |
| **NMA** | 0.31 | 0.34 | 0.35 | 0.37 | 1.00 |

**Table S6** Pearson correlation coefficients between per residue averages of ANISO values. Symmetrical pairwise comparisons were performed between four reference crystals (odd frames).

| **CC ANISO** | **x28** | **x30** | **x33** | **x34** |
| --- | --- | --- | --- | --- |
| **x28** | 1.00 | 0.79 | 0.79 | 0.86 |
| **x30** | 0.79 | 1.00 | 0.69 | 0.72 |
| **x33** | 0.79 | 0.69 | 1.00 | 0.66 |
| **x34** | 0.86 | 0.72 | 0.66 | 1.00 |

**Table S7** Pearson correlation coefficients between per residue averages of ANISO values. Pairwise comparison was performed between four parts of an MD trajectory, the NMA estimate and the experimentally determined ANISO values of the reference crystals (odd frames).

| **CC ANISO** | **x28** | **x30** | **x33** | **x34** |
| --- | --- | --- | --- | --- |
| **MD 1** | 0.36 | 0.25 | 0.35 | 0.34 |
| **MD 2** | 0.39 | 0.24 | 0.35 | 0.35 |
| **MD 3** | 0.32 | 0.23 | 0.23 | 0.31 |
| **MD 4** | 0.38 | 0.24 | 0.31 | 0.34 |
| **NMA** | 0.29 | 0.20 | 0.22 | 0.27 |

Supporting Online Discussion

*Limitations of MD simulations and NMA*

Our original intention was to model the ADP changes that arise upon terahertz irradiation by increasing the amplitude of a selected mode in NMA or increasing the reference temperature in MD simulations. We realized that the ADPs are not modelled as accurately and/or as reproducibly as needed. In two states of the same crystal B_eq_ or ANISO correlate to at least 0.999 and 0.98 (Pearson CC), respectively. Currently, it appears unrealistic to expect that level of correlation between a crystal structure and a simulation or even between two simulations performed with identical parameters. Instead, we switched focus and compared the absolute values of experimental and simulated ADPs. B_eq_ metrics were better estimated by MD, but the level of anisotropy was closer to experimental levels when predicted by NMA. Compared to MD simulations, the confidence interval for some residues even incorporated the mean experimental values (Figure 4B). This indicates that superposition of modes may be a good approach for predicting atom positional distributions and that the relative amplitudes of the modes are not far from realistic. On the other hand, comparison with experimental data indicates that the absolute amplitude of modes and the direction of mode vectors are not accurate. Also, we considered correlation against two massively simplified metrics of ADPs (B_eq_ and ANISO) averaged over multiple atoms only. Unsurprisingly, all NMA modeled atoms form a completely separate cluster from experimental results, when the clustering is performed together (not shown).

Collective eigenvectors derived from NMA (of full atom models or simpler elastic networks (*10*)) are often extremely extrapolated to describe conformational changes. These protein conformations are not expected to fluctuate harmonically. The expectation is especially low at the predicted (terahertz) frequencies due to the large amplitudes involved. The success and suitability of NMA for this purpose may stem from their ability to map the paths of least resistance of an interconnected structure. Nevertheless, for predicting optical phonon modes, a theory that is more fundamental may be needed. Analysis of experimental ADPs revealed intercalating clusters of bonded atoms, which indicates that it is unlikely that a coarse-grained simulation describes physically realistic protein dynamics on the sub-ns timescales. Simplification of MD simulations may be based on experimentally identified (long-range) clusters instead, if synchrony of local modes is the cause of similarity.

*Postulated connection between ADPs and bonding*

Protein atoms are always embedded in a covalent bonding environment, which direct the fluctuation of the atoms in certain directions relative to their neighbors. Atoms in proteins are almost always anisotropic and observing isotropic atoms is extremely improbable.(*11*) To maintain the alignment of these directional fluctuations, one can trivially displace two or more identical chemical groups with the exact same orientation. As an alternative possibility, one can chose a pivot atom in the chemical group, rotate the chemical group by 180º around any eigenvectors of the pivot atom’s ADP without compromising the alignment with the other fluctuating atoms of the same type. This could also explain why only a subset of atoms have a similar ADP when comparing two similar chemical groups: the atoms around the pivot lose the alignment unless they have the exact same ADP as the pivot atom. Analogously, the formation of protein complexes and protein crystals might be mediated by a subset of aligned atoms sparsely distributed in the protein scaffolds. The pre-orientation of the chemical building blocks resulting from the emerging alignment assembly may dramatically reduce the need for random trials during protein folding. The chemical groups (side chains) are held together by the polypeptide chain, which by itself is prone to alignment.

The connection between ADPs and bonding is illustrated in Figure S4 and S5. Figure S4A shows that the main chain carbonyl carbon atoms cluster together in terahertz irradiated crystals. In Figure S4B it is apparent that these atoms are distributed throughout the protein scaffold, but we focus our attention here on the evolutionary conserved region 194-197. As Figure S5 shows the carbonyl groups are parallel in residues Asp-194 and Gly-196 and antiparallel in residues Ser-195 and Gly-197. The carbonyl carbon atoms serve as the pivot point of the amide groups and their ADPs or postulated dominant oscillations are invariant to the rotation. The ADPs of surrounding atoms either passively follow the rotation of coordinating bonds or align with other atoms in the structure. A particularly clear example is the ADP of the elongated Gly-197 Cα atom for which the longest eigenvector is perpendicular to the longest eigenvector in the Gly-196 Cα atom in line with the 90º rotation of the C-Cα bond. As shown previously, the Cβ atom of Ser-195 residue belong to another lattice together with Cβ atom of His-57. The intercalated lattice of pivot atoms may create the long-range geometric stability of folded protein structures.

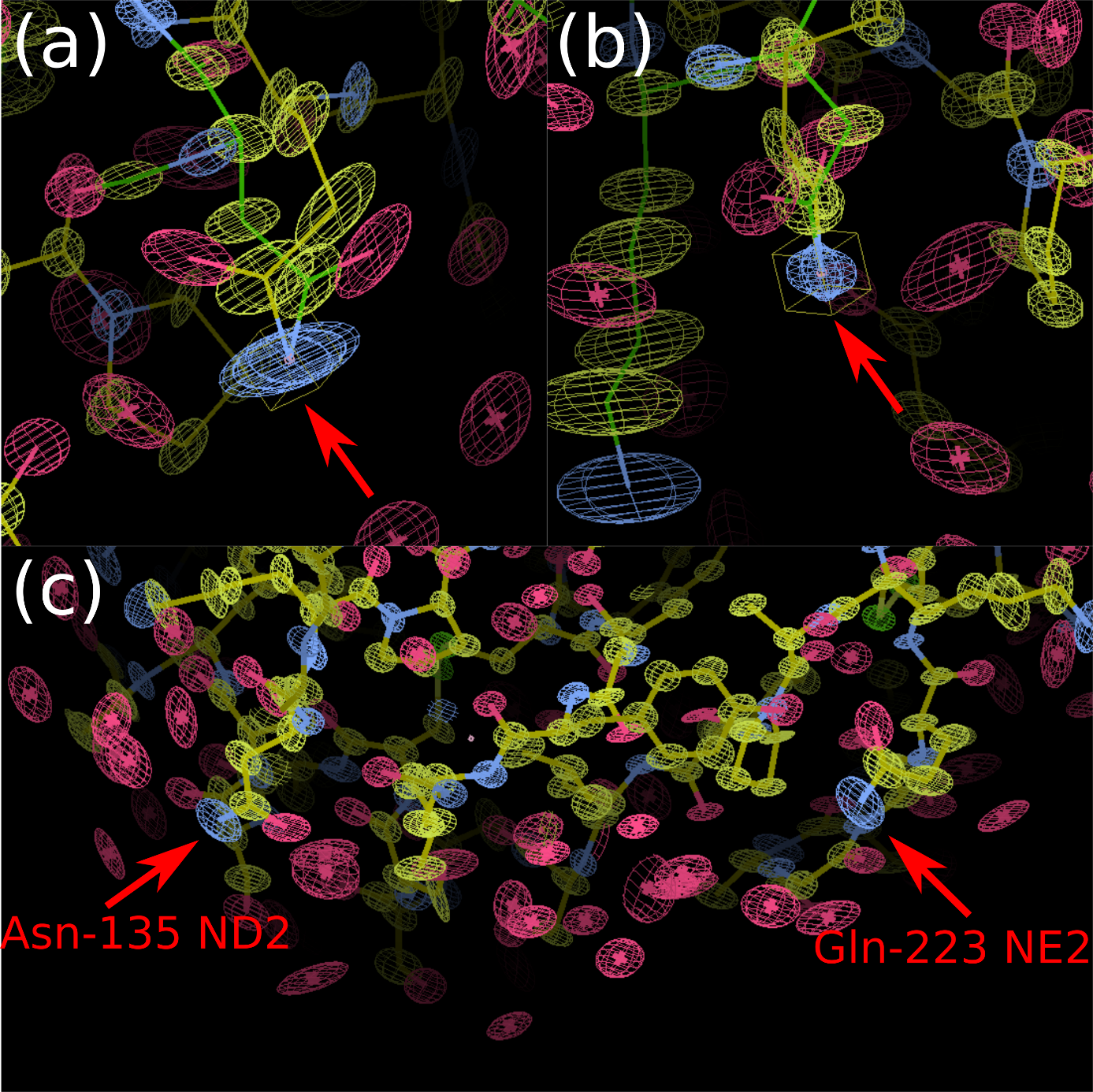

**Figure S2** Comparison of ADP ellipsoids of the side chain amide nitrogen atoms of Gln-135 and Asn-223. *Yellow*, *red*, *blue* and *green* ellipsoids correspond to carbon, oxygen, nitrogen and sulfur atoms, respectively. (a) The two ellipsoids are translated to the identical center point. Structures with y*ellow* and *green* bonds correspond to residues Gln-135 and Asn-233, respectively. (b) 90º rotated view of the same alignment. (c) View of the two atoms as they appear in the trypsin structure.

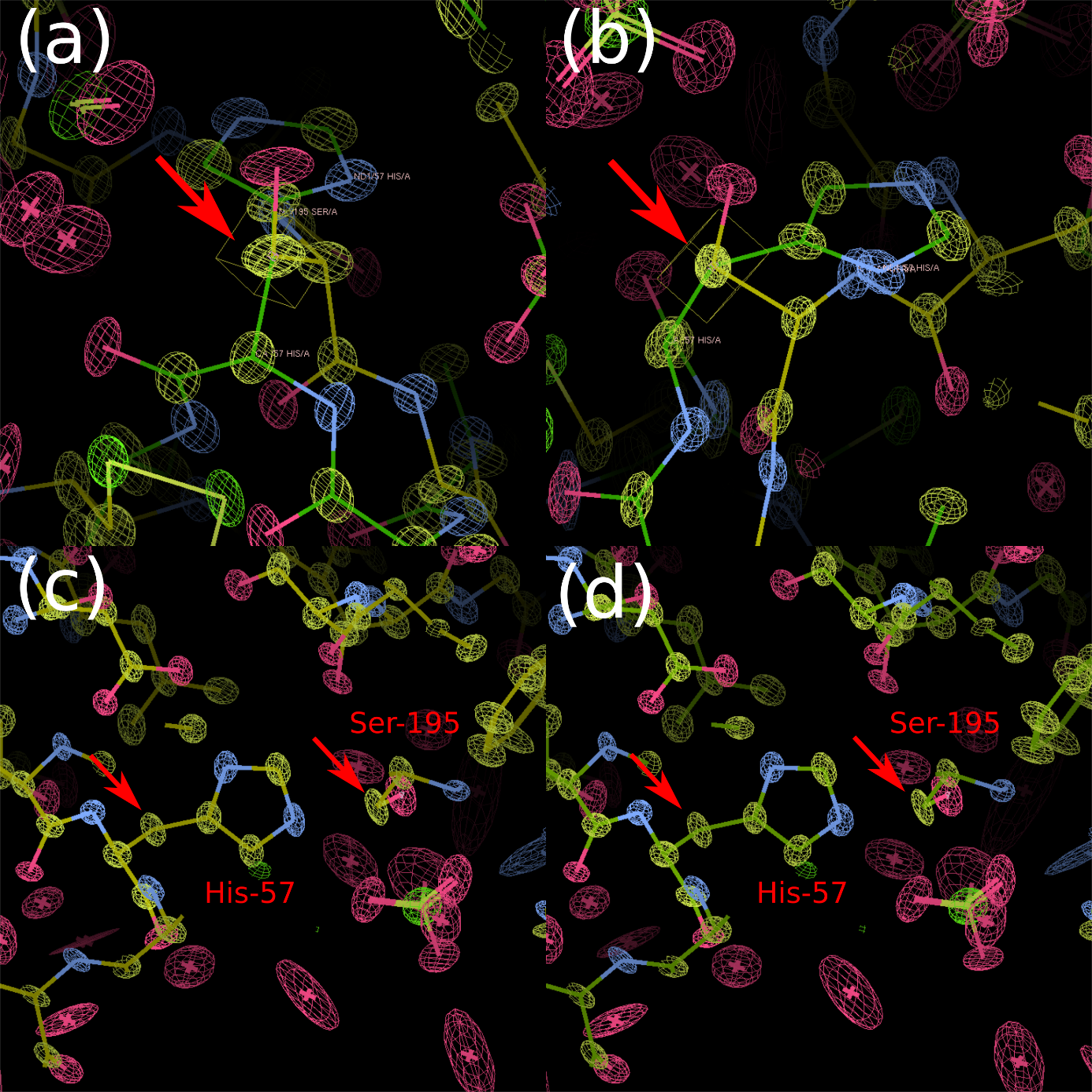

**Figure S3** Comparison of ADP ellipsoids of the Cβ atoms of Ser-195 and His-57. (a) Overlay of the two ellipsoids by translating the center to the same point. *Yellow*, *red*, *blue and green* ellipsoids correspond to carbon, oxygen, nitrogen and sulfur atoms, respectively. *Yellow* and *green* bonds represent to residues Ser-195 and His-57, respectively. (b) The same overlay rotated by 90º. (c) View of the two atoms as they appear in the trypsin structure in the THz on state. (d) The same view of the THz off state.

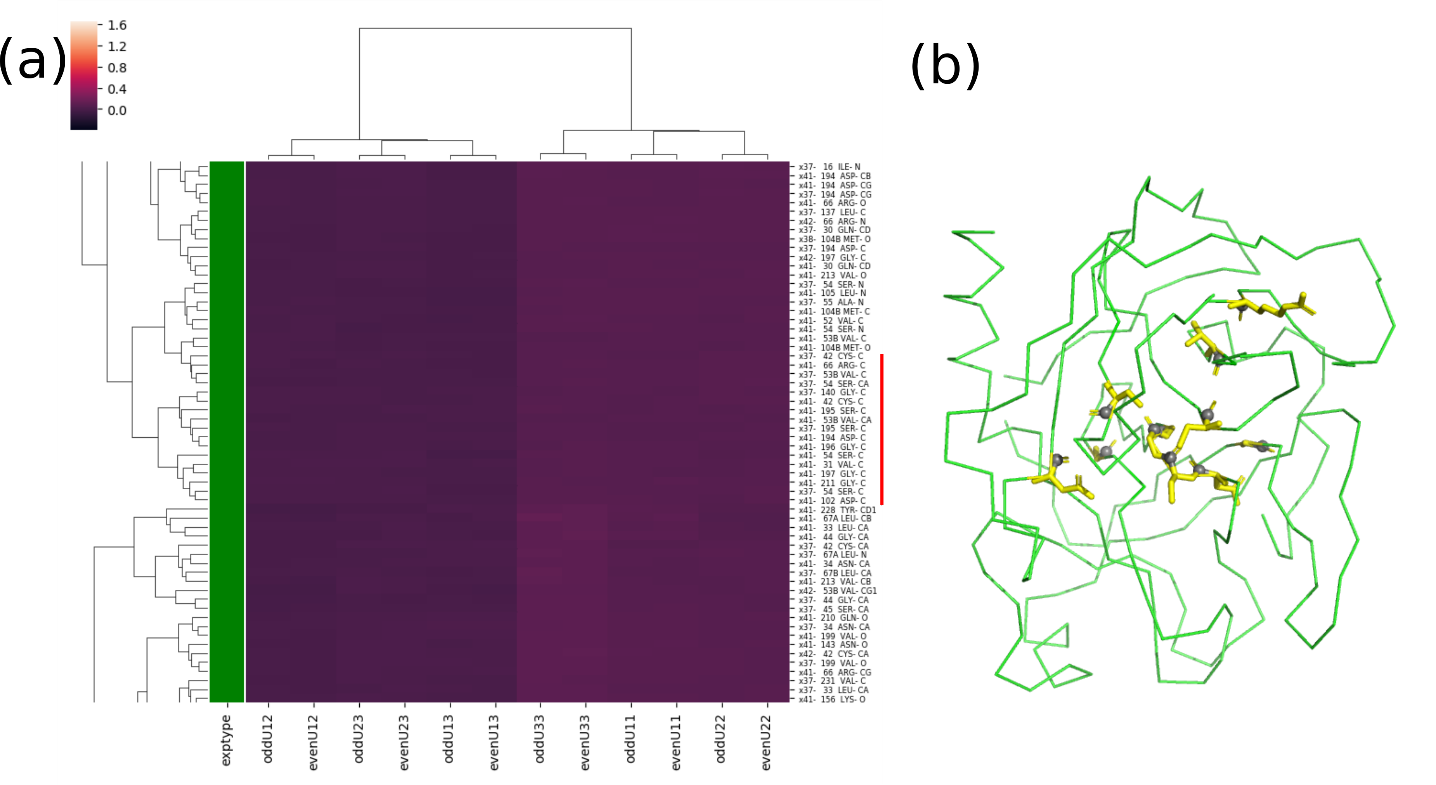

**Figure S4** (a) Clustering of main chain carbonyl carbon atoms. The cluster mainly contains atoms from crystal x41 with some atoms from crystal x37 repeatedly clustering together. Both crystals were irradiated with terahertz radiation. The atoms belong to evolutionary conserved amino acid residues. Asp-102 and Ser-195 are part of the catalytic triad, Asp-194 forms interaction with Ile-16 when the zymogen is activated in serine proteases. The DSGG motif (194-197) is a central signature of chymotrypsin-like serine proteases ([DE]-S-G-[GS]). (b) Spatial distribution of clustered carbonyl carbon atoms are indicated with *gray* balls and the rest of the amino acid residues is marked by *yellow* sticks. The Cα trace of trypsin is depicted as a *green* ribbon.

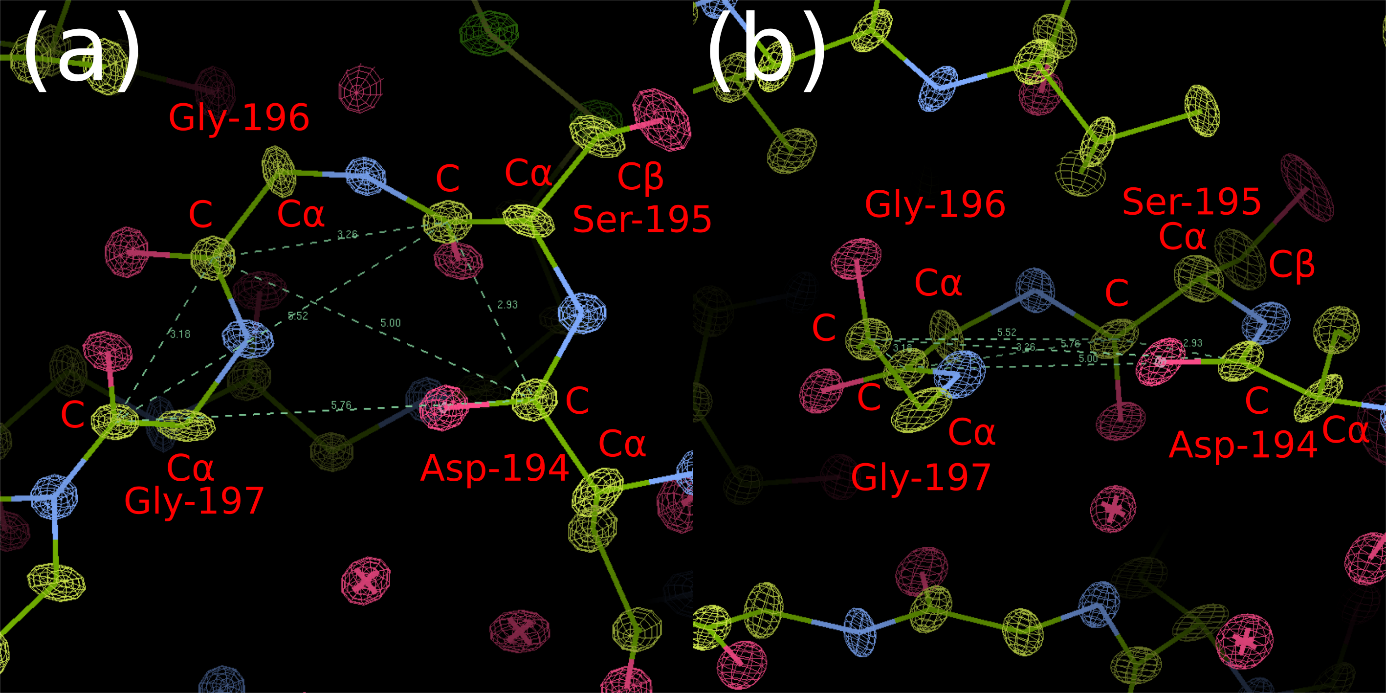

**Figure S5** Magnified view of the lattice structure in the central DSGG motif (residues 194-197). *Yellow*, *red* and *blue* ellipsoids correspond to carbon, oxygen and nitrogen atoms, respectively. (a) View perpendicular to the plane defined by the carbonyl carbon atoms in this motif. (b) View parallel with the plane.

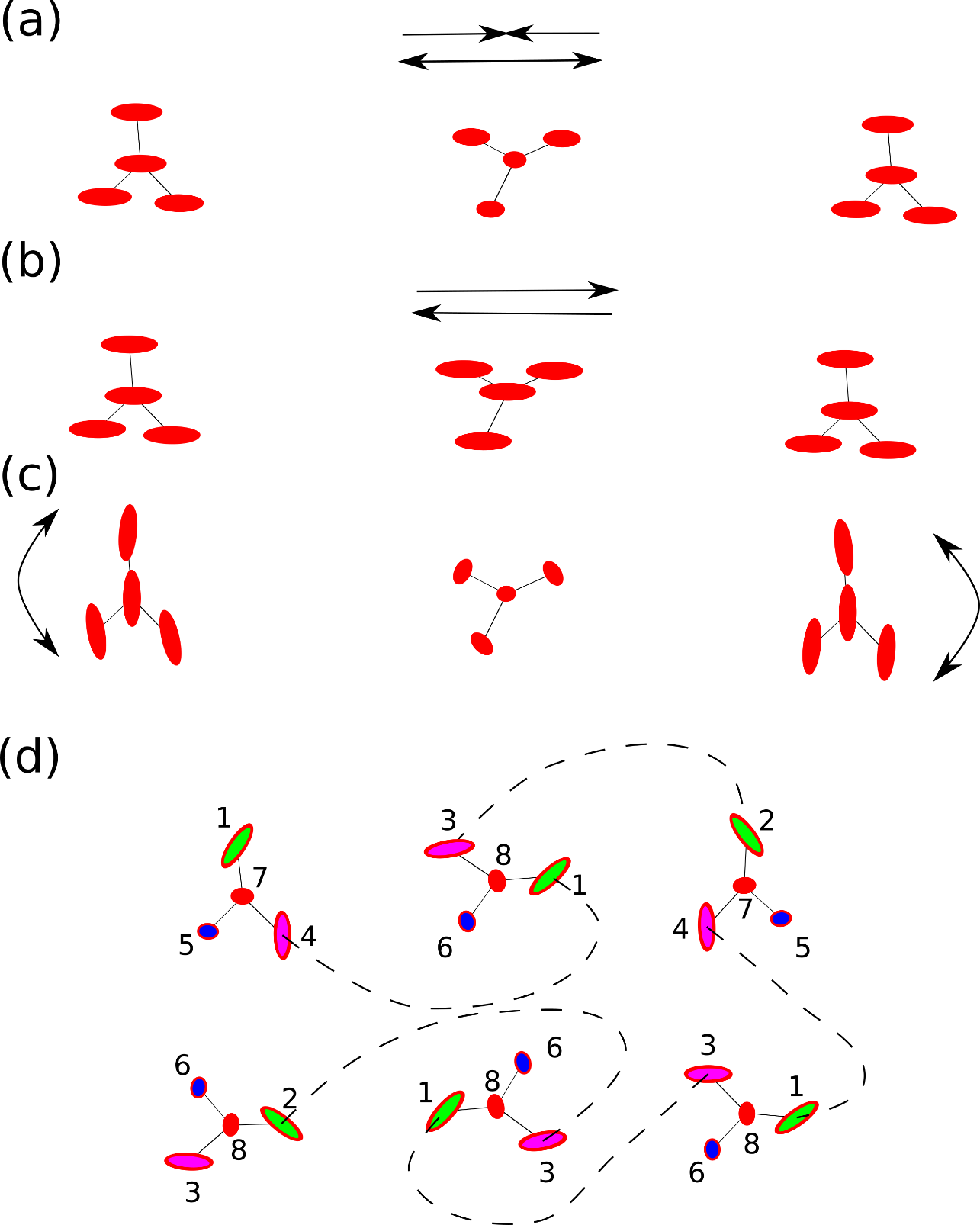

**Figure S6** Different vibrational mechanisms lead to distinct patterns of atomic displacements in chemical groups. (a) Delocalized elastic stretch could occur in a protein that has a positive and negatively charged patch connected by a “neutral” elastic region. Alternating electric fields would periodically stretch and compress the entire protein. Elastic network simulations describe this type of dynamics. (b) Rigid body translation/rattling could occur in alternating electric fields if the entire protein has a net positive or negative charge. (c) Rigid body rotation or delocalized elastic bending. The common features in the delocalized vibrational mechanisms in (a-c) is that adjacent atoms have similar displacements. Although the entire protein may preferentially orient through these mechanisms, this would not contribute to the orientation of internal components. (d) In our model, partial charges are distributed alternately in the protein and they are specific to the atom type. The colors represent different atom types. Adjacent atoms with different partial charges react differently to alternating electric fields, therefore the atomic displacements will also depend on the type of atom. Symmetry operations permit the alignment of similar atoms types. Therefore, the same orientation of the entire chemical group (translational symmetry) is not necessary. This can give rise to multiple lattices that are vibrating in concert at the same time. Atoms with the same number share similar displacement and they would cluster together when subjected to Ward’s method.
